# Feature similarity: a sensitive method to capture the functional interaction of brain regions and networks to support flexible behavior

**DOI:** 10.1101/2024.11.13.623324

**Authors:** Xiuyi Wang, Baihan Lyu, Katya Krieger-Redwood, Nicholas E. Souter, Golia Shafiei, Nan Lin, Jonathan Smallwood, Elizabeth Jefferies, Yi Du

## Abstract

The brain is a dynamic system where complex behaviours emerge from interactions across distributed regions. Accurately linking brain function to cognition requires tools that are sensitive to these dynamics. We introduce a novel technique, Feature Similarity (FS), which represents a conceptual advancement in measuring brain region interactions by integrating multiple features across various measures, moving beyond traditional methods that focus on a single measure. Our results show that FS can capture functional brain organisation: regions within the same functional network have greater FS compared to those in different networks, and FS also identifies the principal gradient that spans from unimodal to transmodal cortices. FS demonstrated greater sensitivity to task modulation than traditional functional connectivity (FC) and 46 out of 49 statistical measures of pairwise interactions (SPIs). Specifically, FS reveals interaction patterns missed by FC and most SPIs, such as a double dissociation in Dorsal Attention Network: greater interaction with Visual network during working memory tasks and greater interaction with default mode network during long-term memory tasks. These findings position FS as a powerful tool for capturing task-specific brain network dynamics and advancing our understanding of cognitive flexibility. Using FS to reanalyse existing fMRI data could resolve inconsistencies, lead to novel insights, and prompt revisions of existing theories or the development of new theoretical frameworks.

## 1. Introduction

A central aim of cognitive neuroscience is to understand how brain regions interact to support flexible behaviour. The increasing availability of large-scale datasets, where participants perform multiple cognitive tasks, provides a valuable opportunity to address this question. However, existing methods typically focus on a single feature of interaction, such as co-variability, failing to capture the multidimensional nature of brain dynamics [1]. As a result, these methods struggle to reliably track task-dependent changes in network interactions. Since the choice of interaction metric fundamentally shapes our understanding of the brain’s functional organization, there is a critical need for a more comprehensive approach that integrates multiple features to capture context-dependent functional interactions.

Current neuroimaging methods typically measure specific aspects of functional interactions but overlook their complex, multidimensional nature. Many rely on temporal correlation, assessing neural synchrony but with notable limitations. For example, within-subject functional connectivity (FC) cannot distinguish stimulus-driven from intrinsic correlations, reducing task sensitivity [2], while inter-subject FC isolates stimulus-driven effects but requires identical stimuli across participants, limiting its applicability [2]. Psycho-physiological interactions assess task-dependent connectivity but rely on predefined seed regions and assume a linear relationship with task conditions, restricting flexibility [3]. Dynamic causal modelling infers directed interactions but requires a priori model specification, assumes stationarity, and is computationally limited to small networks [4]. Recently, many statistics of pairwise interaction (SPIs) from fields like Earth system and finance have been applied to brain data [1,5], yet they still focus on single interaction features (e.g., causality, co-variability) rather than integrating multiple dimensions. Consequently, these unidimensional metrics often yield inconsistent or contradictory findings. For instance, frontoparietal control network (FPCN) exhibits negative FC with default mode network (DMN) [6], suggesting opposing functions, yet both share similar timescales [7], implying shared functional properties. Such inconsistencies underscore the limitations of unidimensional approaches and highlight the need for a multidimensional method sensitive to task-dependent modulations.

A time series can be characterized by numerous features, each capturing an interpretable structure [8,9] that corresponds to brain functions. Gaussianity belongs to distributional properties, capturing statistical characteristics such as normality and outliers, which influence higher-order measures of variability and predictability. Autocorrelation belongs to temporal dependencies, capturing how past values influence future values, essential for assessing dynamic coupling between brain regions. Entropy belongs to predictability, capturing signal uncertainty, where higher entropy indicates greater unpredictability. Each feature corresponds to distinct brain functions, as indicated by observations that certain brain regions exhibit specific features, equipping them for different roles, while regions with similar features tend to support parallel functions, regardless of their FC. For instance, early visual areas and DMN occupy opposing ends of the timescale hierarchy: early visual areas exhibit rapid autocorrelation decay, whereas DMN regions show gradual decay [2,7,10–12]. This timescale difference underlies their functional roles: neural representations in early visual areas are minimally influenced by prior knowledge, while the DMN is heavily shaped by it [13]. Similarly, transmodal regions such as FPCN and DMN, though often showing negative FC [6], share long timescales[7] and process similar information over longer periods (i.e., maintaining task goals) [13,14]. These findings suggest that multidimensional features reveal functional similarities beyond those captured by FC or SPIs, which focus on single aspects of neural interaction.

Features dynamically shift depending on cognitive tasks [15,16], indicating their potential to depict variations in network interactions. For example, timescale of neural responses varied across tasks [15,16]: during auditory story comprehension, transmodal regions exhibit a shortened autocorrelation window, while unimodal regions maintain it, whereas motor and working memory tasks exhibit the opposite effect [16]. These shifts reflect the brain’s adaptive reconfiguration, as seen in the domain-general FPCN, which exhibits greater FS with the dorsal attention network (DAN) during working memory tasks and FS with the memory control network during long-term memory tasks [6]. These observations motivated us to develop feature similarity (FS): a hypothesis-free, data-driven method designed to capture how interaction patterns vary across tasks. FS extracts a wide range of interpretable features from time-series data, calculates FS between brain regions, and evaluates how FS varies across tasks (Fig. 1). Since tasks impose specific cognitive demands, the brain’s functional organization is modulated at multiple levels. Temporal correlation (as captured by FC) and other unidimensional features (as captured by SPIs) may remain stable, yet the underlying signal complexity, variability, and temporal autocorrelation may exhibit greater task sensitivity. This multidimensional analysis enables FS to detect subtle brain dynamics that may be missed by existing unidimensional methods.

**Fig. 1.**
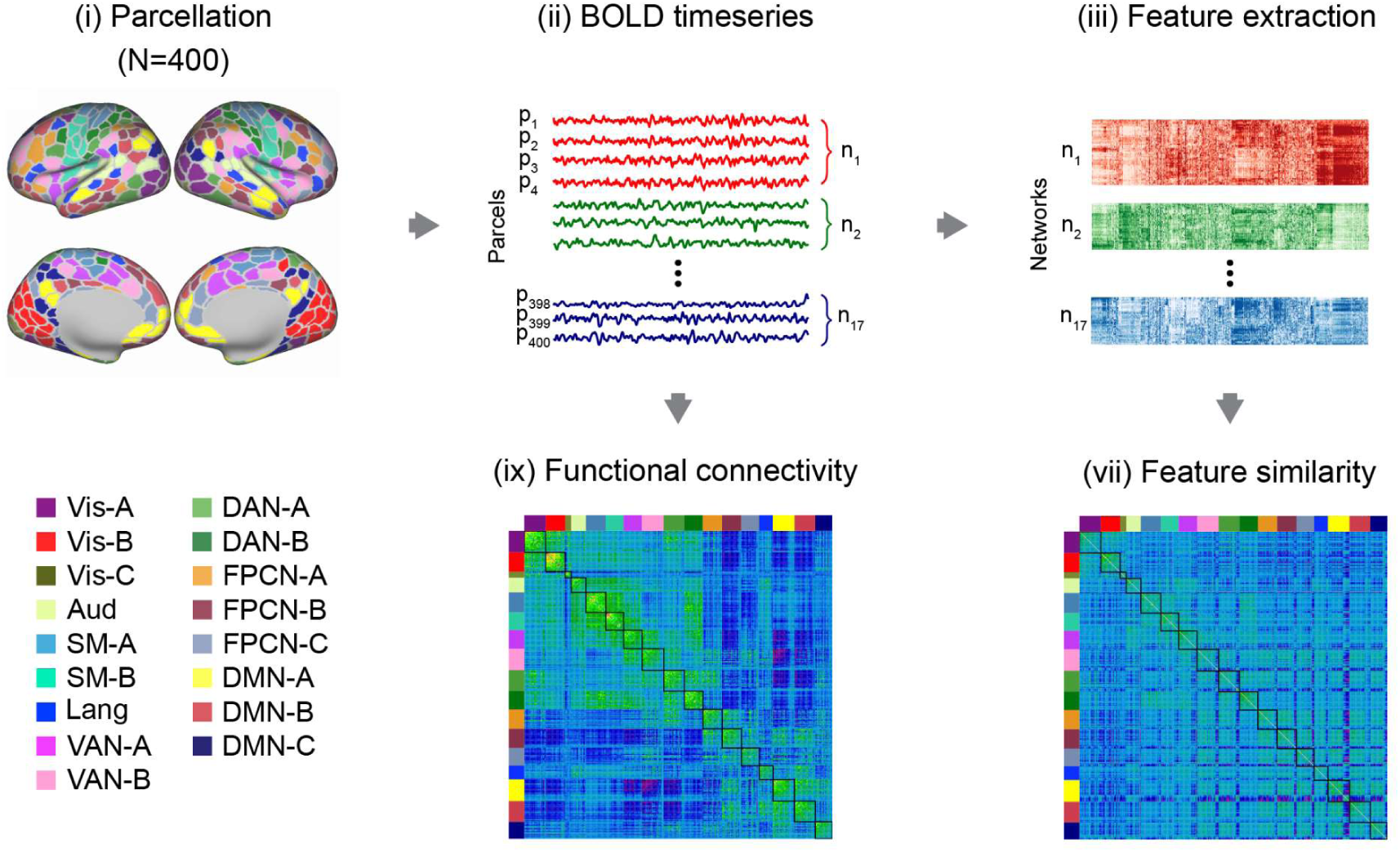
The workflow of the FS and FC analysis. (I) Individual-specific parcellation divided the whole brain into 400 parcels across 17 networks [19]. (II) Average time series of each parcel. (III) Extraction of features of time series for each parcel. (IV) Pearson correlation coefficients of the extracted features represent the pairwise feature similarity between all possible combinations of brain parcels. (V) Functional connectivity involved calculating Pearson correlation coefficients between the time-series of parcels. (Vis = Visual, Aud = Auditory, SM = Sensory-motor, DAN = Dorsal attention network, VAN = Ventral attention network, FPCN = Fronto-parietal control network, Lang = Language, DMN = Default mode network; p1–p400 represent the 400 parcels, and n1–n17 represent the 17 networks in the analysis).

**Fig. 2.**
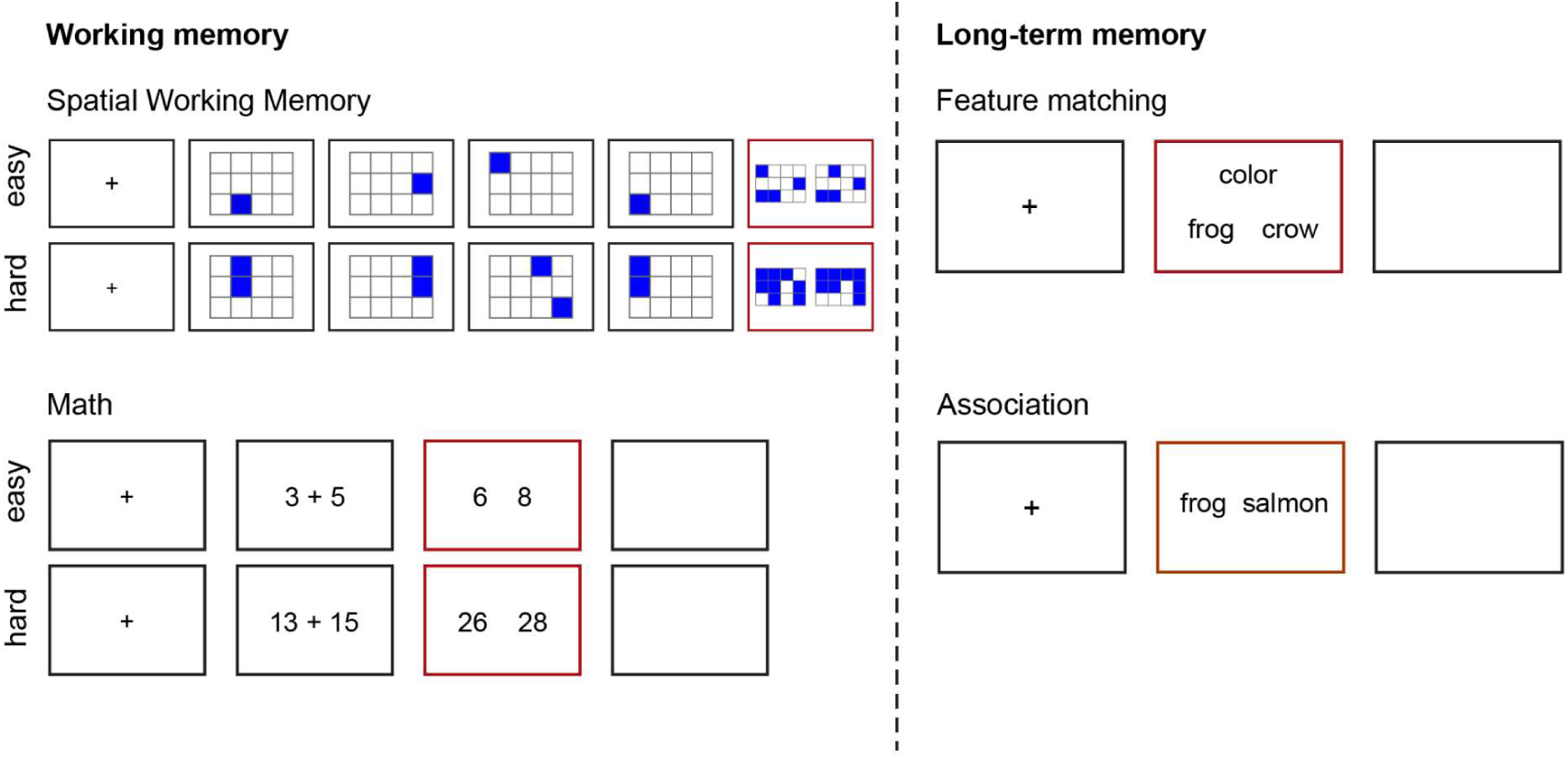
The experimental design. To tap working memory, we included two tasks: a spatial working memory task required participants to keep track of sequentially presented locations, while math decisions involved maintaining and manipulating numbers which rely more on working memory. To tap long-term memory, we included two tasks that required controlled retrieval of knowledge; a semantic feature matching task required participants to match probe and target concepts according to a particular semantic feature (color or shape), while a semantic association task involved deciding if pairs of words were linked in meaning. Response periods are indicated by a red box.

To evaluate FS’s sensitivity in capturing task-dependent network interactions, we examined the functional relationship between DAN and DMN. Traditionally, DAN, which directs external attention, has been considered antagonistic to DMN, which supports internally directed cognition such as memory retrieval and semantic processing [17]. This opposition is reflected in their typical anticorrelation and DAN’s strong FC with the Visual network, reinforcing its role in sensory-driven attention [18,19]. However, recent evidence challenges this strict dichotomy. Using multiple metrics, DAN has been shown to be topographically positioned between the Visual network and DMN, allowing it to flexibly interact with both based on task demands [16,20]. Moreover, DMN-DAN interactions are integrated into broader network dynamics involving the FPCN [18], which shares properties with both DMN and DAN and plays a pivotal role in flexible cognition by bridging external attention and internal processing. FPCN cooperates with DMN via the memory control network during long-term memory tasks but not during working memory tasks [6], suggesting that DAN, too, may align more with DMN in long-term memory tasks, shifting from external sensory attention to internally guided retrieval. Our results demonstrate that FS captures these task-driven shifts more effectively than FC and 46 out of 49 SPIs, revealing a double dissociation: DAN interacts more with the Visual network during working memory tasks and with DMN during long-term memory tasks—a distinction that FC and most SPIs fail to detect. These findings highlight FS’s unique ability to uncover nuanced, task-dependent brain dynamics, establishing it as a powerful tool for studying cognitive flexibility in brain networks.

## 2. Results

The results are divided into three sections. (i) First, we took an existing individualized parcellation of the cortex to identify the parcels of each network and examined whether FS is equally capable of capturing the network structure as FC does. As expected, regions belonging to the same intrinsic functional networks had greater FS compared to regions in different networks. (ii) Next, we examined whether FS could capture the organizational principles of the cortex as is seen in FC. We found FS captured the principal intrinsic connectivity gradient that separates sensory-motor regions from transmodal areas, as well as the second component that separates somatomotor and auditory from visual cortex, which have been previously described using decompositions of FC. (iii), Finally, we examined whether FS is more sensitive to task-induced changes in brain dynamics than FC and most SPIs by examining how these metrics characterise network interactions across working memory and long-term memory tasks. Specifically, we asked whether FS is superior than FC and SPIs in their capacity to identify differences in the way that DAN interacts with Visual network and DMN during working memory and long-term memory tasks.

The FS method consists of the following steps (see Fig. 1):

Step 1: Extract a diverse set of interpretable features from the time-series data.

Step 2: Calculate FS between brain regions by calculating the Pearson correlation coefficients of the extracted features.

Step 3: Examine how FS varies across different tasks.

### 2.1. Regions belonging to the same intrinsic functional network have greater FS compared to regions in different networks

We first examined whether parcels within the same network exhibited similar features. We selected Kong et al.’ parcellation [19] as it allows for individual-specific parcellations with greater homogeneity [6] while capturing the inherent heterogeneity of large-scale networks like DAN and FPCN. To estimate FC, we computed Pearson correlation coefficients between regional time-series, and for FS, we computed Pearson correlations between regional feature vectors (Fig. 1). Regions were considered similar in terms of features if the features of two regions were significantly correlated [9]. FS captures regions with similar dynamical properties but not necessarily synchronous activity (Fig. 1). For example, while FPCN-A and DMN both have longer timescales, they often exhibit negative FC [6].

As expected, regions within the same functional network showed higher FC compared to those between networks, both at rest (t = 136.09, p < 0.001) and during tasks (spatial working memory, t = 26.33, p < 0.001; math, t = 26.36, p < 0.001; semantic feature matching, t = 70.02, p < 0.001; semantic association, t = 57.60, p < 0.001) (Fig. 3). Similarly, FS was higher within networks compared to between networks, both at rest (t = 91.24, p < 0.001) and across tasks (spatial working memory, t = 18.35, p < 0.001; math, t = 14.51, p < 0.001; semantic feature matching, t = 42.27, p < 0.001; semantic association, t = 38.04, p < 0.001) (Fig. 3). All p-values were FDR-corrected. These findings suggest that FS can capture intrinsic network structure traditionally measured by FC.

**Fig. 3.**
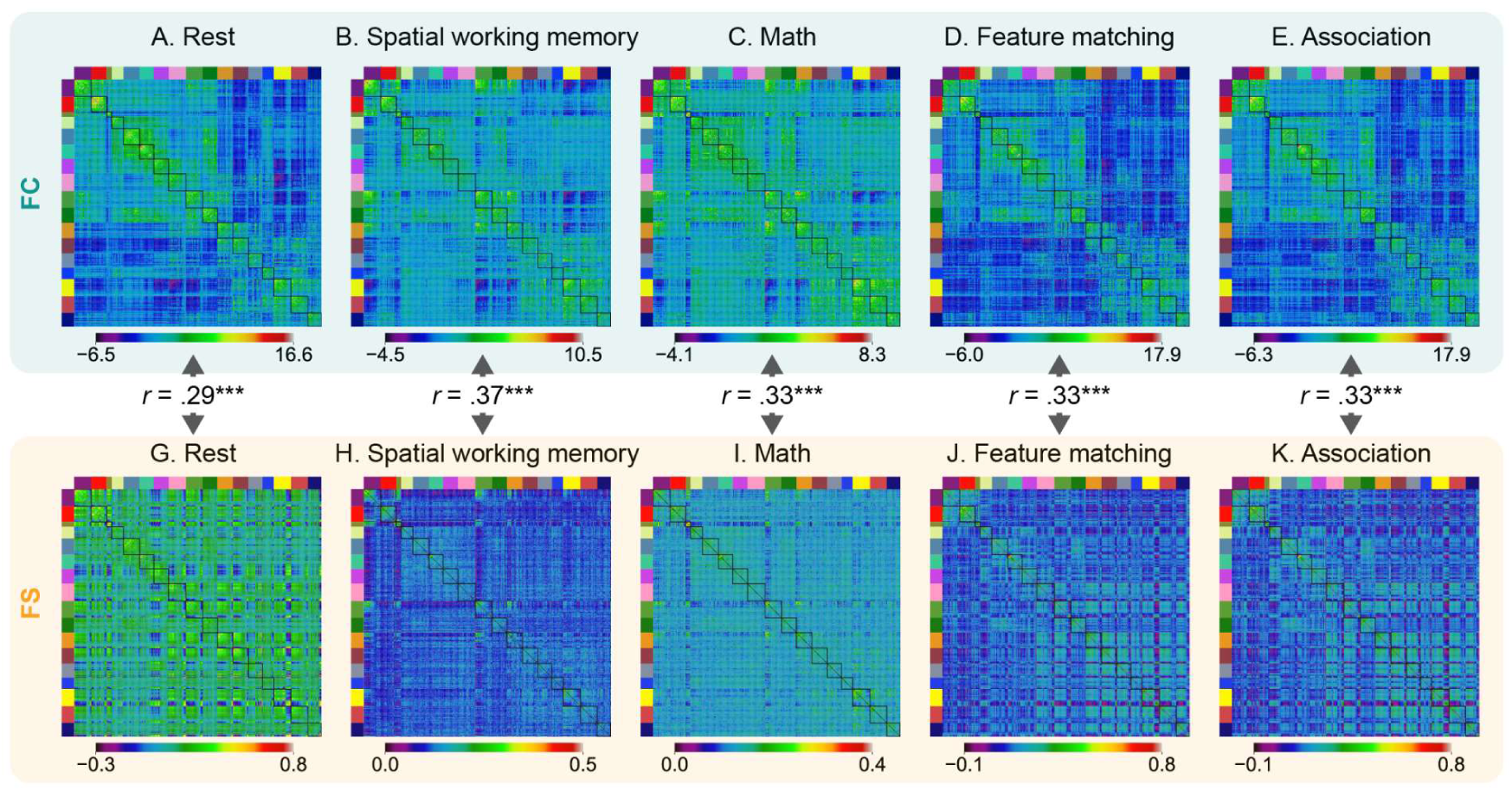
The FC matrix and FS matrix at rest and for each task. Regions within the same intrinsic functional network exhibited greater FC and FS than regions in different networks at rest and for each task. FC and FS showed positive but weak correlations at rest and for each task. (***: p < 0.0001).

We also assessed the similarity between FC and FS by correlating the FC and FS matrices at rest and during tasks. We observed weak but significant positive correlations between FC and FS as indicated by arrows (Fig.3) at rest (r = 0.29, p = 0) and during tasks (spatial working memory, r = 0.37, p = 0; math, r = 0.33, p = 0; semantic feature matching, r = 0.33, p = 0; semantic association, r = 0.33, p = 0). All p-values were FWE-corrected using maximum r values (Method 4.5.4). The positive correlations suggest that regions with similar time-series features exhibited coherent spontaneous fluctuations. However, the largest correlation was 0.37, indicating that these measures also capture different information.

### 2.2. FS reveals unique organizational principles beyond FC

Next, we investigated whether FS and FC captured similar or different components of connectivity. Firstly, we performed dimension reduction analysis (diffusion embedding) [20] on the resting state FC matrix for the HCP dataset (Fig. 3A). We focused on the three components with the largest eigenvalues, as they explain 28.02% variance in total and have clear interpretation (see Fig. 4I for scree plot). Consistent with previous studies [20–23], the first component explaining 12.75% of the variance corresponded to the principal gradient described by Margulies et al. (2016) [20]. This component separated sensory-motor regions (shown in purple-blue in Fig. 4A) from transmodal areas (shown in red-white in Fig. 4A). The second component explained 11.29% of the variance and separated somatomotor and auditory cortex (shown in purple-blue in Fig. 4B) from visual cortex (shown in red-white in Fig. 4B). The third component explained 3.98% of the variance and separated FPCN regions (shown in purple-blue in Fig. 4C) from DMN regions (shown in red-white in Fig. 4C).

**Fig. 4.**
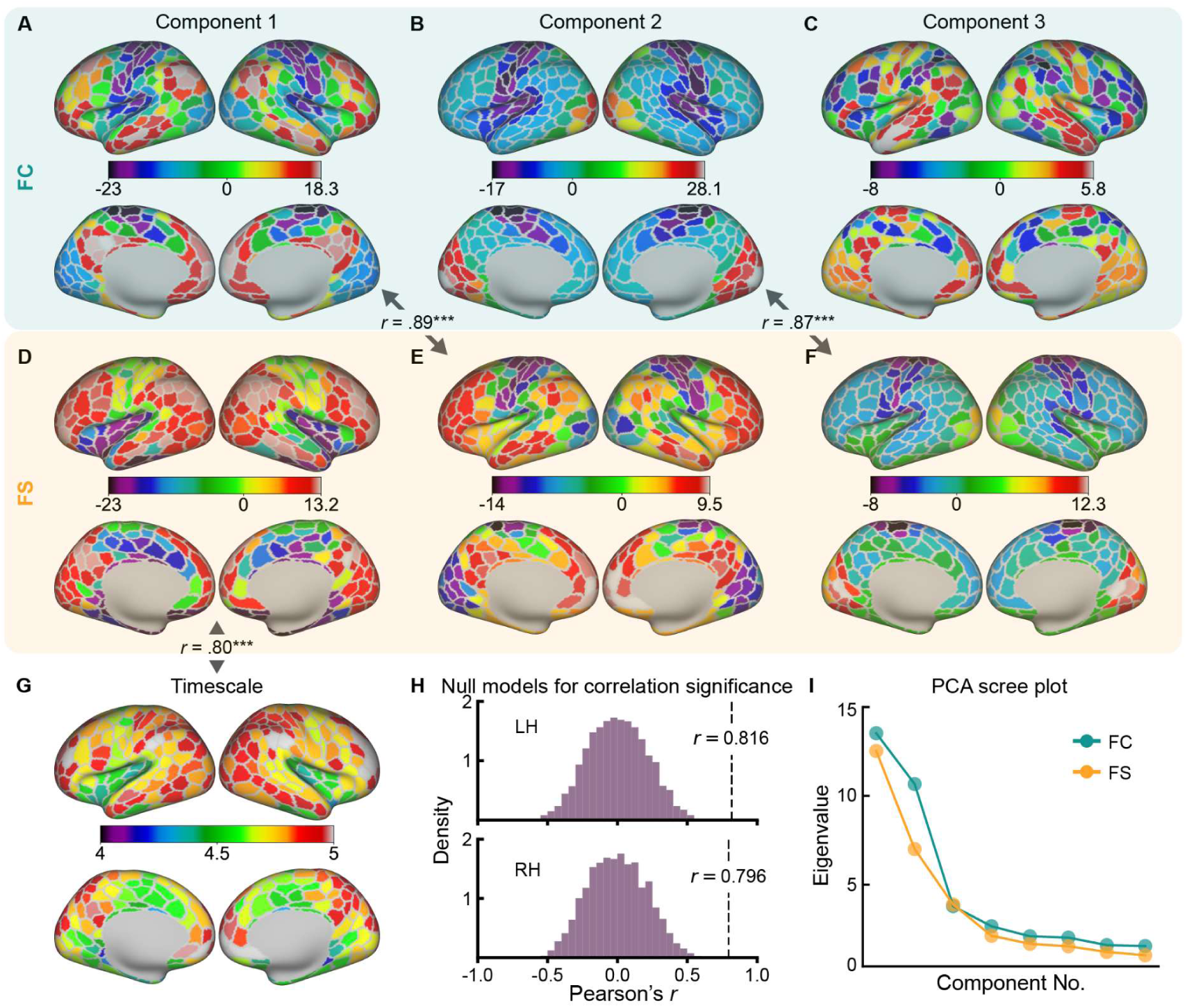
The top three principal components of FC and FS. FC and FS captured two similar principal components and one distinct component. The first principal component of FC (A) corresponds to the second principal component of FS (E), as indicated by the arrows denoting a strong correlation (r = 0.89, p = 0, corrected for spatial autocorrelation using spin permutation), which separates sensory-motor regions from transmodal areas. Similarly, the second principal component of FC (B) corresponds to the third principal component of FS (F), with a strong correlation (r = 0.87, p = 0, corrected for spatial autocorrelation using spin permutation), separating somatomotor and auditory cortex from visual cortex. These findings suggest that FS captures similar organizational information to FC. However, the third principal component of FC (C), which separates FPCN regions from DMN regions, was not captured by FS. The first principal component of FS (D), which corresponds to the intrinsic timescale gradient (G), was not captured by FC. (G) The intrinsic timescale map shows short timescales in insula and cingulate cortex, and long timescales in angular gyrus, posterior cingulate cortex, and frontal pole. (H) The first principal component of FS was significantly correlated with the intrinsic timescale gradient map in both the left (r = 0.82, p = 0) and right (r = 0.80, p = 0) hemispheres (corrected for spatial autocorrelation using spin permutation). The histograms illustrate the null model distributions. (I) Scree plots showing the eigenvalues for the top eight principal components of FS (orange) and FC (blue).

Similarly, we performed dimension reduction analysis on the resting state FS matrix for the HCP dataset (Fig. 3F), using the procedures above. The three components with the largest eigenvalues explained 23.23% of the variance (see Fig. 4I for scree plot). FS captured two components that were similar to the results for FC, and one not uncovered by FC (Fig. 4). The first principal component of FS explained 12.19% of the variance and had a distinct spatial distribution from the first three components of FC: it did not show significant correlations with any of the components of FC (p > 0.05; Fig. 4D). The next two components of FS were seen in FC data (Fig. 4). The second component of FS (explaining 7.11% of the variance; Fig. 4E) aligned with the principal gradient of FC (Fig. 4A), separating sensory-motor from transmodal cortex, with a strong correlation between components (r = 0.89, p = 0, spin permutation corrected). The third component of FS (explaining the 3.94% variance; Fig. 4F) corresponded to the second component of FC (Fig. 4B), separating somatomotor and auditory cortex from visual cortex, with a strong correlation between components (r = 0.87, p = 0, spin permutation corrected). These findings further confirm that FS captures meaningful information, i.e., organizational principle.

To gain insight into the first component of FS, we investigated its correlation with the map of intrinsic timescale, which reflects the temporal duration of ongoing inputs that the brain can process. The first component resembles the distribution of intrinsic timescale reported by Raut et al. (2020) [7], which is longest in transmodal regions [15]. We therefore estimated intrinsic timescale for each parcel by measuring the decay of the temporal autocorrelation function and quantifying the time taken for the autocorrelation function to reach a threshold of r= 0.5, which is half of the full width at half maximum. Higher values indicate longer processing times for ongoing inputs. Our results align with Raut et al. (2020) [7], showing short timescales in insula and cingulate cortex, and long timescales in angular gyrus, posterior cingulate cortex, and frontal pole. (Fig. 4G). We found that the first principal component, which explained the most variance in FS, was significantly correlated with the intrinsic timescale map (left hemisphere: r = 0.82, p = 0; right hemisphere: r = 0.80, p = 0; Fig. 4H, corrected for spatial autocorrelation using spin permutation), indicating that FS can capture meaningful information not captured by traditional FC, such as the intrinsic timescale of the brain’s response.

### 2.3. FS showed greater variation across tasks than FC

The above analyses showed that FS captured similar information to FC but also additional dimensions of cortical organisation. Next, we examined whether FS was more sensitive to task modulation than FC. FC measures the temporal correlation between time-series from different brain regions, focusing on the synchrony of neural activity. In contrast, FS extracts a comprehensive set of features from the time-series, such as entropy, stationarity, and autocorrelation, providing a multidimensional profile of the brain’s activity. Given that tasks impose specific cognitive demands and engage distinct neural processes, the brain’s functional organization undergoes modulation at multiple levels. Temporal correlation (as captured by FC) may remain stable under certain conditions, but the underlying features of the time-series, such as signal complexity, variability, and temporal autocorrelation, may exhibit more sensitivity to these changes. This multidimensional analysis may allow FS to detect subtle changes in brain dynamics during tasks that may be missed by FC.

We computed the Pearson correlation coefficients of the task mean FC matrices for every possible combination pair of tasks. For example, we calculated the Pearson correlation coefficients of the FC matrices of the spatial working memory task (Fig. 3B) and math task (Fig. 3C). Similarly, we calculated the Pearson correlation coefficients of the FS matrices of these two tasks (Fig. 3G and 3H). Then we directly compared these two r values. We found correlations for FC were consistently higher than those for FS for every task pair (Fig. 5A; p < 0.05; see Table 1 for statistics), suggesting that FS may be more sensitive to task differences than FC.

**Fig. 5.**
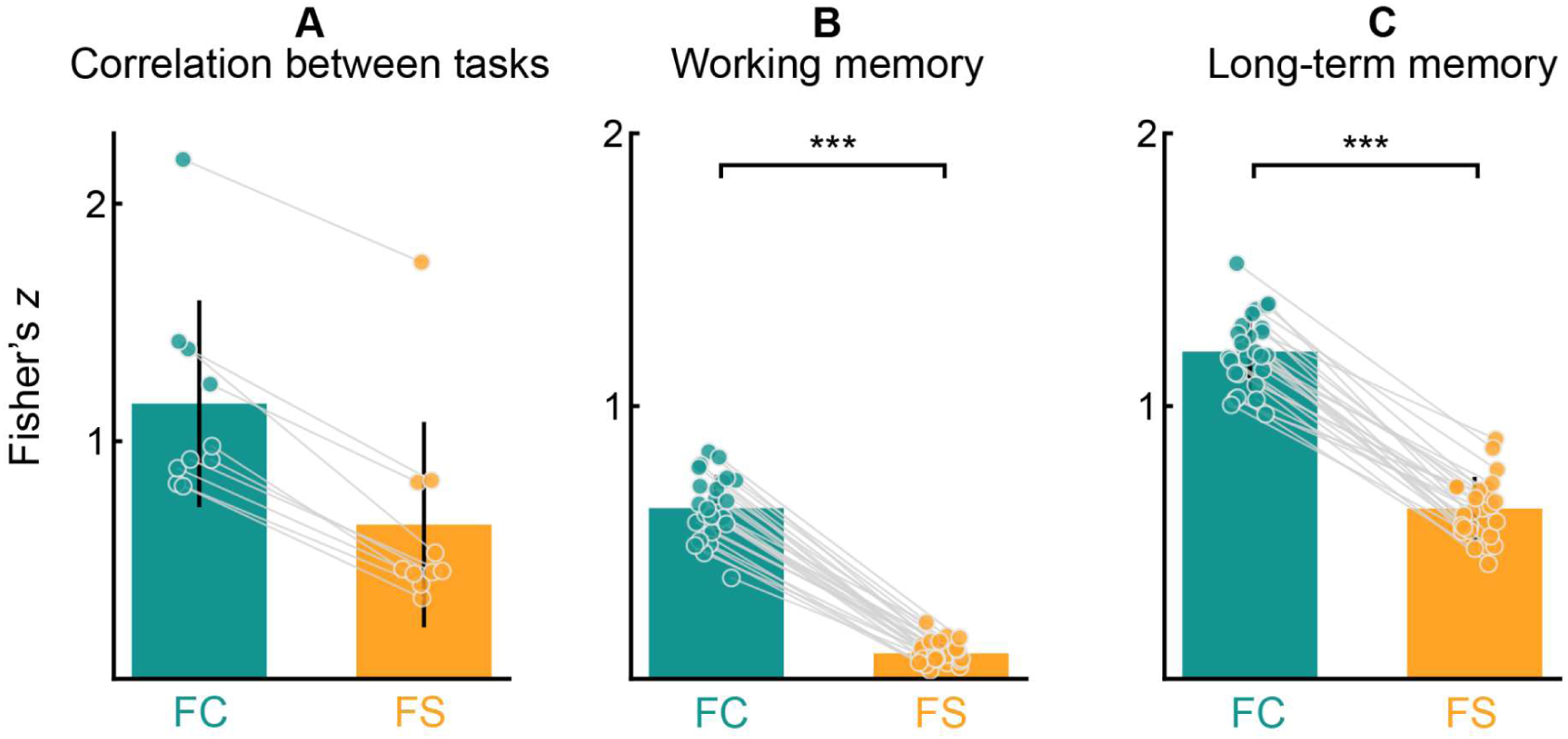
FS showed smaller correlations across tasks than FC. A – The group mean FC matrices showed higher correlations across tasks compared to FS matrices. Each dot represents the correlation coefficient between two tasks, with 10 possible combinations among the four tasks. B – The correlations of FC matrices for spatial working memory and math task were greater than the correlations of FS matrices. Each dot represents the correlation coefficient between the two task-specific matrices of one participant. C – The correlations of FC matrices for semantic feature matching and semantic association task were significantly greater than the correlations of FS matrices for these two tasks. Each dot represents the correlation coefficient between the two task-specific matrices of one participant.

**Table 1.** FS showed greater variation across tasks than FC.

| Task 1 | Task 2 | r (FC) | r (FS) | z | p |
| --- | --- | --- | --- | --- | --- |
| Rest | Spatial working memory | 0.68 | 0.33 | 96.31 | 0 |
| Rest | Math | 0.67 | 0.38 | 82.64 | 0 |
| Rest | Feature matching | 0.86 | 0.68 | 82.48 | 0 |
| Rest | Association | 0.88 | 0.68 | 110.49 | 0 |
| Spatial working memory | Math | 0.75 | 0.43 | 103.38 | 0 |
| Spatial working memory | Feature matching | 0.73 | 0.42 | 94.50 | 0 |
| Spatial working memory | Association | 0.71 | 0.42 | 88.63 | 0 |
| Math | Feature matching | 0.75 | 0.43 | 103.38 | 0 |
| Math | Association | 0.73 | 0.43 | 93.72 | 0 |
| Feature matching | Association | 0.98 | 0.94 | 86.35 | 0 |

The task sensitivity of FS was further confirmed when we compared the correlations of FS with those of FC between two non-semantic tasks for each participant. Since the same participants completed both the spatial working memory task and the math task, we calculated the Pearson correlation coefficients between the FC matrix of the spatial working memory task and that of math task for each participant and transformed the resulting r values to z values using Fisher’s transformation. Similarly, we also calculated the Fisher’s z values between the FS matrices of the two non-semantic tasks for each participant. Finally, we compared the z values of FC with those of FS by conducting paired t-tests. We found that the correlations of FC were greater than the correlations of FS for the non-semantic tasks (Fig. 5B; t = 26.58, p < 0.001). We then conducted the same analysis for the two semantic tasks, semantic feature matching and association, and observed the same pattern (i.e., stronger correlation across tasks for FC than for FS; Fig. 5C, t = 23.29, p < 0.001). The smaller correlation of FS is not because the data of FS are noisier. By contrast, FS carries meaningful task information, as FS matrices correctly classify task labels for non-semantic tasks (accuracy = 0.83, p = 0) and semantic tasks (accuracy = 0.78, p = 0), both significantly above chance level (0.5) (see SM 2.1 for detailed results). These findings suggest that FS is more sensitive to task modulation than FC, as FS takes into account multiple aspects of time-series similarity, whereas FC reflects only one.

### 2.4. FS captured network interaction patterns across tasks not captured by FC

After showing that FS was generally more sensitive to task modulation than FC, we examined whether FS could capture the varying network interaction patterns across tasks that missed by FC. We tested the interaction difference between DAN and Visual network versus DAN and DMN, to test the hypothesis that DAN is more similar to Visual network in non-semantic tasks, which rely more on visual features and working memory, and more similar to DMN in semantic tasks, which rely more on long-term memory. We expected that this pattern would be seen more readily in FS, since this metric was more sensitive than FC to task demands above.

Since the tasks were presented visually in the current study, we selected DAN-A [24], which is typically engaged together with Visual network and showed greater feature similarity with Visual network than other attention networks (DAN-B, Ventral attention network) in a classification analysis (Fig. S1). We compared FC and FS between DAN-A and Visual network versus DAN-A and DMN across tasks, focusing on differences in the strength of these connections. For simplicity, we combined the three visual subnetworks into a single visual network and the three default mode subnetworks into a single DMN. While connectivity differences did not vary across tasks when assessed with FC (Fig. 6, p > 0.05, FWE corrected), there was a difference between non-semantic and semantic tasks for FS (p < 0.001; FWE corrected). DAN-A always showed greater FC with Visual network than with DMN at rest (Fig. 6, t = 53.68, p < 0.001) and for each task (Fig. 6, spatial working memory, t = 16.19, p < 0.001; math, t = 11.73, p < 0.001; semantic feature matching, t = 17.19, p < 0.001; semantic association, t = 14.95, p < 0.001), consistent with previous studies. However, DAN-A showed greater FS with Visual network than with DMN at rest (Fig. 5, t = 10.58, p < 0.001) and in non-semantic tasks (Fig. 5, spatial working memory, t = 3.55, p = 0.004 ; math, t = 3.19, p = 0.003), but showed the opposite pattern in semantic tasks (Fig. 5, semantic feature matching, t = -2.86, p = 0.008; semantic association, t = -2.64, p = 0.01). These findings suggest that FS is more sensitive to dynamic interaction patterns between networks that reflect differing cognitive demands.

**Fig. 6.**
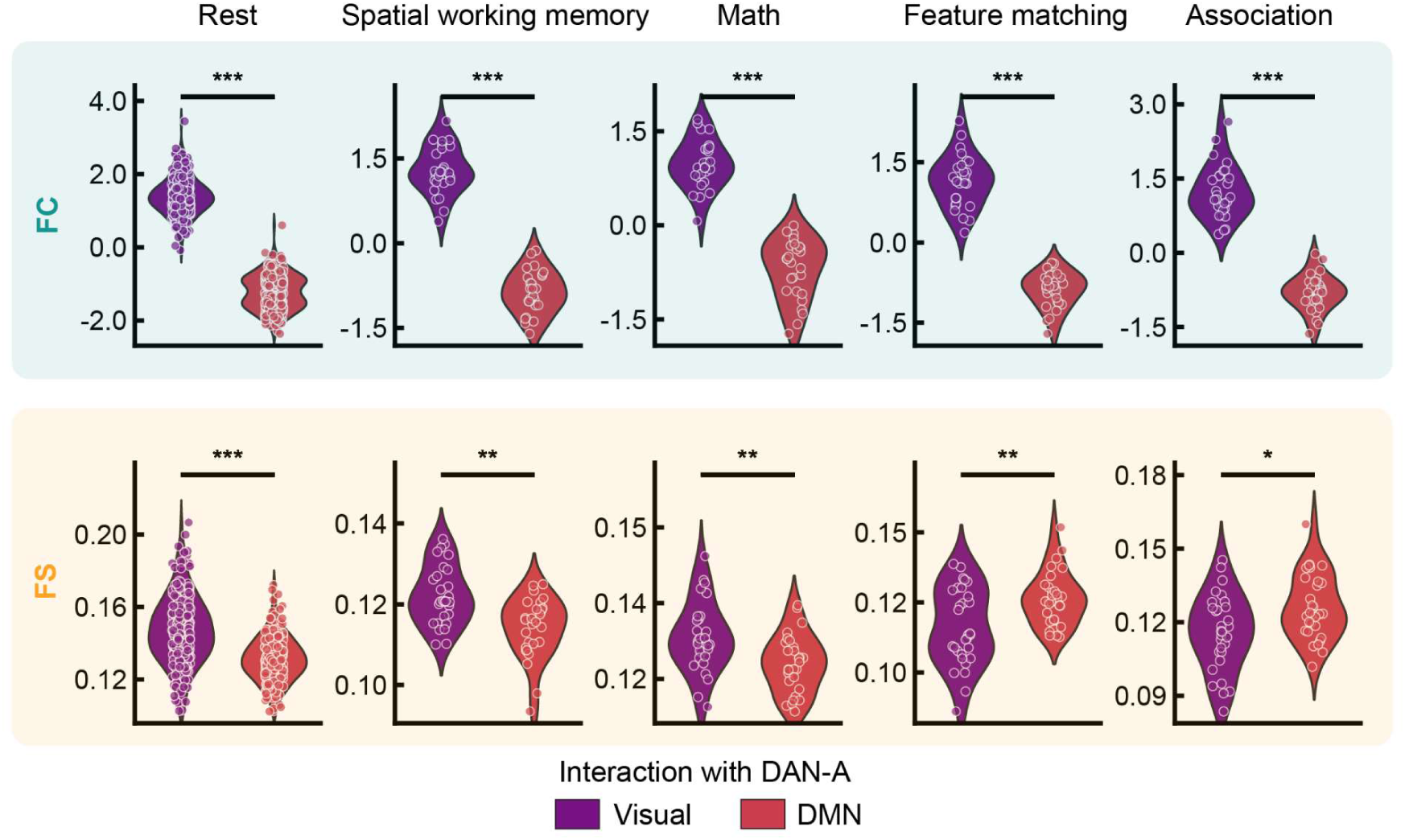
FS captured varying interaction patterns for DAN-A across tasks that missed by FC. Top panel: DAN-A always showed greater FC with Visual network than with DMN at rest and for each task and there was no difference across tasks. Bottom panel: The difference between DAN-A and Visual network versus DAN-A and DMN in the non-semantic tasks was greater than in semantic tasks. DAN-A showed greater FS with Visual network than with DMN at rest and for the non-semantic tasks, but showed the opposite pattern in the semantic tasks that relied on long-term memory supported by DMN.

The task sensitivity of FS was further confirmed when we examined the interaction differences between the domain general control network (FPCN-A) and Visual network versus FPCN-A and DMN. Interaction differences for FPCN-A across tasks were only captured by FS and not by FC (Fig. S2; See SM 2.3 for detailed information). There was no difference in FC across tasks for FPCN-A (p > 0.05, FWE corrected), with FPCN-A always showing greater FC with Visual network than with DMN at rest (t = 2.39, p = 0.02) and for each task (spatial working memory, t = 11.36, p < 0.001; math, t = 11.44, p < 0.001; semantic feature matching, t = 6.50, p < 0.001; semantic association, t = 3.62, p = 0.001). However, FS revealed different patterns across tasks, with a greater difference between FPCN-A and Visual network compared with FPCN-A and DMN for the non-semantic than semantic tasks (p < 0.001; FWE corrected). Specifically, FPCN-A showed similar FS with Visual network and DMN at rest (t = 1.39, p = 0.17) and in the math task (t = -0.73, p = 0.47) but showed greater FS with DMN than with Visual network in the spatial working memory task (t = -4.97, p < 0.001) and semantic tasks (semantic feature matching, t = -9.12, p < 0.001; semantic association, t = -8.25, p < 0.001). These findings support that FS is more sensitive to task modulation than FC.

### 2.5. FS captured network interaction patterns across tasks not captured by most SPIs

While FC (Pearson correlation) remains the dominant method for measuring brain network interactions in fMRI, recent studies have shown that other SPIs, previously applied in different fields, can also be used to measure neural interaction since some of them can classify brain states and conditions in fMRI data [1]. To further assess FS’s sensitivity relative to these SPIs, we initially selected all 67 SPIs that significantly classified states in the fMRI film dataset and then refined this set to 49 SPIs with reasonable computational requirements (<5 hours per run per task per subject). These SPIs derived from 20 interaction measures across six categories: 1) Basic Methods, such as precision, which quantifies pairwise associations while controlling for the effects of other time series; 2) Distance-Based Similarity, which measures statistical similarity or independence based on pairwise distances between bivariate observations; 3) Causal Inference, which aims to infer directed relationships; 4) Information-Theoretic Measures, such as mutual information which quantifies the total dependency between two variables; 5) Spectral Measures, commonly computed in the frequency or time-frequency domain. Examples include spectral coherence magnitude, which quantifies the alignment of frequency components in phase and amplitude; and 6) Miscellaneous Methods: this category includes techniques such as linear model fits, which estimate relationships through regression, and other statistical tools that quantify pairwise interactions but do not fit into the above categories (see Methods 4.5.11 and Supplementary Material 1.3.3 for full details). We examined whether the double dissociation observed by FS can be replicated by each SPI by comparing the interaction matrices generated by each SPI across tasks, focusing on key networks (DAN-A, Visual, and DMN). Specifically, we conducted paired t-tests for each task to assess differences between network pairs to examine whether DAN-A showed greater interaction with Visual network during the working memory task but with DMN during the semantic tasks.

We found that 46 out of the 49 SPIs did not fully replicate the double dissociation observed with FS. Of these, 21 methods lacked task sensitivity: 20 showed consistently stronger interactions between the DAN and Visual networks across all tasks, while 1 showed the opposite pattern (Fig. 7A). Examples include covariance from the Basic Category and Longest common subsequence from the Distance Similarity Category (Fig. 7C). 10 SPIs demonstrated limited sensitivity, each detecting only a single dissociation. Among these, 9 showed stronger DAN-Visual interactions during working memory tasks but failed to capture the reverse pattern in semantic tasks, such as Mutual Information from the Information Theory Category with a Gaussian kernel (Fig. 7D) (Fig. 7A). In contrast, 1 SPI exhibited the opposite dissociation, revealing stronger DAN-DMN interactions during semantic tasks while failing to detect the reverse pattern in working memory tasks.

**Figure 7.**
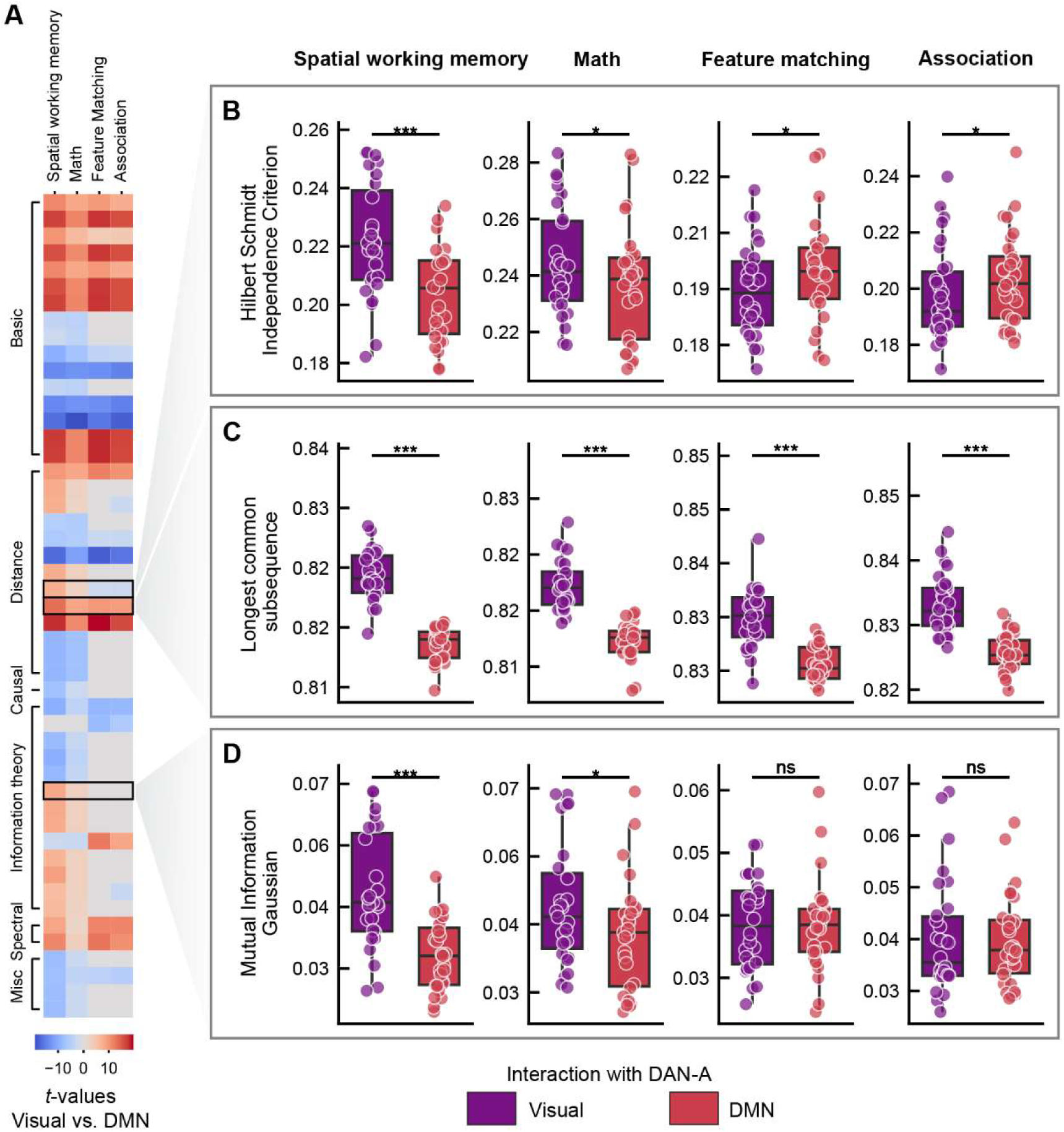
Task-dependent interactions of DAN-A with Visual (purple) and DMN (red) networks as measured by 49 SPIs, illustrating varying levels of task sensitivity. (A) Heatmap showing the task-dependent interactions of DAN-A with Visual and DMN networks across all 49 SPIs. Colours represent t-values comparing interactions between networks, with blue indicating stronger Visual network interactions and red indicating stronger DMN interactions. (B) Hilbert-Schmidt Independence Criterion from the Distance Similarity category, a representative SPI that successfully reveals the double dissociation, showing greater DAN-A-Visual interaction during working memory and greater DAN-A-DMN interaction during semantic tasks. (C) Longest common subsequence from the Distance Similarity Category, a representative SPI that lacks task sensitivity, consistently showing greater DAN-A-Visual interaction across tasks. (D) Mutual Information from the Information Theory category, a representative SPI that reveals a single dissociation, showing greater DAN-A-Visual interaction during non-semantic tasks but failing to reveal the reverse pattern for semantic tasks. Significance levels are indicated (*p < 0.05, **p < 0.01, ***p < 0.001, ns = not significant). These examples illustrate the variability in task sensitivity across SPIs, underscoring the unique sensitivity of FS to task modulation.

Only 1 SPI, Hilbert-Schmidt Independence Criterion from the Distance Similarity Category, fully replicated the double dissociation, capturing task-specific interactions of DAN with the Visual network during working memory tasks and the DMN during semantic tasks (Fig. 7B). 2 SPIs: Distance Correlation from the Distance Similarity category and time-lagged mutual information from the Information Theory category partially replicated the double dissociation, failing to show significantly stronger DAN-DMN interactions than DAN-Visual during the feature matching task (FDR-corrected p = 0.07).

The remaining SPIs included 14 methods showing a reverse single dissociation and 1 method displaying a reverse double dissociation, but these results lacked clear interpretability, further highlighting the challenges of using SPIs to detect task-specific patterns. Additionally, we observed that SPIs with different parameter settings could yield inconsistent results. For instance, precision with estimator EmpiricalCovariance revealed a reverse single dissociation, while the version with the estimator ShrunkCovariance showed no task sensitivity. Full statistical results of each SPI are provided in Supplementary Table 2. These findings underscore the difficulty of selecting optimal SPIs and parameters for specific tasks.

The failure of most SPIs to replicate the double dissociation observed with FS underscores the superior sensitivity of FS in detecting nuanced brain network dynamics. Among the 49 SPIs tested, only a few (3 SPIs) successfully replicated the double dissociation, further validating FS’s findings and robustness. This dual observation highlights FS’s unique ability to capture task-specific interactions that are often missed by other methods, offering critical insights into cognitive flexibility and functional reconfiguration.

## 3. Discussion

In this study, we introduce FS as a robust method for unveiling network structures, brain topography, and task-induced modulations. Our validation of FS is supported by two key observations: (1) Intrinsic network consistency: regions within the same intrinsic functional network exhibit greater FS compared to regions across distinct networks, and (2) Functional hierarchy mapping: FS captures the principal gradient that organizes neural systems along a continuum from unimodal (sensory-motor regions) to transmodal (high-order cognitive areas) cortices. Building upon previous research, our findings demonstrate that FS possesses heightened sensitivity to task modulation, revealing unique interaction patterns undetected by traditional FC and 46 out of 49 SPIs. For instance, DAN-A exhibits greater FS with Visual network than DMN during working memory tasks, and the opposite pattern during long-term memory tasks. This double dissociation suggests a functional shift in DAN-A towards DMN during semantic tasks that engage long-term memory processes. In contrast, FC fails to detect this distinction, consistently showing greater connectivity between DAN-A and Visual network across both task types. While 10 out of 49 SPIs reveal single dissociations (such as DAN showing greater cooperation with the Visual network during non-semantic tasks), only 3 out of 49 SPIs capture the double dissociation identified by FS. These discrepancies underscore FS’s superior ability to detect nuanced, task-dependent brain dynamics, offering compelling evidence of its advantage over existing methods. The enhanced task sensitivity of FS is further corroborated by its detection of interaction patterns of FPCN-A. Collectively, our findings position FS as a promising tool for elucidating complex network interaction patterns and advancing our understanding of brain function.

Our findings demonstrate that FS is a more effective method for analysing interaction patterns across cognitive domains compared to FC and other SPIs, owing to its greater sensitivity to task effects. FS can be utilized to reanalyse existing datasets, potentially resolving inconsistencies in prior research and leading to novel insights. The brain operates as an intricate information processing and transfer system, necessitating coordination among networks supporting diverse functions to facilitate complex cognitive processes, such as memory retrieval [25]. Examining the interaction patterns of brain networks offers a deeper understanding of this dynamic organization. While FC has traditionally been used to examine these patterns, its limited sensitivity to task modulation may have constrained our understanding of the flexible nature of network cooperation. Although numerous SPIs have been proposed to measure interactions [1,5], they often focus on single aspects and remain less sensitive to task modulation than FS. We suggest that reanalyzing fMRI data using FS could yield novel insights, resolve inconsistencies, and inspire revisions to existing theories or the development of new frameworks.

FS represents a conceptual advancement in measuring brain region interactions by integrating multiple features across various dimensions, moving beyond the traditional focus on covariance in FC. While FC has been instrumental in understanding neural connections, it primarily captures linear relationships, potentially overlooking complex, multidimensional interactions. Recent developments in SPIs have introduced methods that assess specific aspects of neural interactions; however, each tends to concentrate on a single dimension, limiting their scope. In contrast, FS encompasses a broader spectrum of features, enabling a more comprehensive analysis of neural dynamics. This multidimensional approach allows researchers to capture intricate patterns of brain activity that may be missed when relying solely on unidimensional measures. As the field progresses, the specific features incorporated into FS can evolve and expand, enhancing its comprehensiveness. Nonetheless, the fundamental advantage of FS lies in its capacity to provide a holistic assessment of neural interactions, offering a more nuanced understanding of brain function compared to existing methods.

While we advocate the use of FS for reanalysing existing data, this does not imply that FC and other SPIs are without merit. FS serves as a complementary measure to FC and other SPIs, which remains valuable for comprehending the general functional architecture of the brain, for example, as shaped by neuroanatomy. FC is particularly adept at revealing coupling patterns of intrinsic activity. For instance, DMN regions exhibit greater FC with each other at rest and during various conditions and this functional similarity is thought to be at least partly related to their anatomical location at a maximum distance from primary sensory-motor cortex [20] and by the strong influence of spatial proximity on connectivity patterns. Yet these observations might underplay the dynamic nature of connectivity patterns and their sensitivity to changes in cognitive state. DMN regions display no ISFC at rest, nor when participants listen to scrambled words or individual words – although they do exhibit ISFC when participants listen to sentences and paragraphs [2]. This observation suggests that FC is predominantly influenced by intrinsic activity, while ISFC is driven by stimulus-related activity during states of coherent semantic cognition. Since FC, ISFC and FS convey distinct information, combining these methods can offer a comprehensive understanding of the brain’s functionality.

Although this study highlights the increased sensitivity of FS to task modulation, the specific features that change across tasks have yet to be determined. It is imperative for future research to identify and interpret features. For instance, we demonstrated that the DAN-A exhibits greater FS with Visual network than DMN in non-semantic tasks, wherein participants focus more on external visual information. Conversely, DAN-A displays greater FS with DMN than Visual network in semantic tasks, where participants rely on long-term memory. Given that approximately 7,000 features were extracted and employed in calculating FS, future research can pinpoint the specific features that vary across these tasks. While our results demonstrate that FS can exhibit greater sensitivity to task modulation than FC in specific datasets and networks, this finding does not imply that feature similarity is universally superior. Therefore, our study should be viewed as an illustration of the potential advantages of FS, rather than a definitive conclusion about its superiority in all contexts. Future investigations should encompass a more extensive array of diverse tasks.

## Conclusion

Our results demonstrate that Feature Similarity (FS) offers a significant methodological advancement in understanding dynamic brain network interactions. While traditional functional connectivity (FC) and other statistics of pairwise interactions (SPIs) methods have provided valuable insights, FS is more sensitive and reliable in capturing subtle changes in network interactions, particularly across different cognitive states. Just as motion tracking software has revolutionized the study of animal behaviour by enabling precise analysis of how animals interact in different social contexts, we argue that FS represents a similar breakthrough in neuroimaging—enhancing our ability to assess brain function and cognitive flexibility with greater precision. While experimental designs and neuroimaging hardware continue to improve, FS provides a powerful tool for reanalysing existing fMRI datasets, resolving inconsistencies, and advancing our theoretical understanding of cognitive flexibility. Ultimately, FS could become an essential tool for exploring the adaptive nature of brain network dynamics.

## 4. Materials and Methods

This study included three datasets, one publicly available dataset – the Human Connectome Project (HCP), and two task fMRI datasets collected at the University of York, UK.

We analysed the resting state functional MRI (rsfMRI) data of 245 unrelated participants who completed all four resting state scans from the S900 release of HCP dataset to investigate the functional hierarchy, FC, and FS patterns. In addition, to compare whether FC and FS can capture varying interaction patterns to the same extent, we used two types of tasks (tapping working memory and long-term memory retrieval) which are associated with distinct neurocognitive modes [26] To tap working memory, participants completed easy and hard spatial working memory and arithmetic tasks designed to localise domain general control regions [27–29], which required information to be maintained and manipulated. They also completed two semantic tasks tapping knowledge in long-term memory, a semantic feature matching task that involved linking probe and target concepts (presented as written words) based on colour or shape, and a semantic association task in which participants decided if pairs of words were semantically associated or not [6,23]. These datasets have been previously analysed [6,23].

### 4.1. Participants

All participants were right-handed, native English speakers, had normal or corrected-to-normal vision, and had no history of psychiatric or neurological illness. For the HCP dataset, informed consent was obtained, and the study was approved by the Institutional Review Board of Washington University at St. Louis. For the York non-semantic and semantic dataset, the research was approved by the York Neuroimaging Centre and Department of Psychology ethics committees.

We analysed the data of 245 neurologically healthy volunteers (130 males, 115 females), aged 23 – 35 years (mean = 28.21, SD = 3.67), from the HCP dataset [30].

31 neurologically healthy adults (26 females; age: mean ± SD = 20.60 ± 1.68, range: 18 – 25 years) performed spatial working memory and math tasks in York. One participant with incomplete data (only one of two sessions) was removed. A functional run was excluded if (i) mean relative root mean square (RMS) framewise displacement was higher than 0.2 mm, (ii) it had more than 15% percentage of total frames with motion exceeding 0.25 mm, or (iii) a participant’s accuracy on the respective task was three standard deviations below the group mean. If only run of one task was left for a given participant after exclusion, the data from this task were removed for this participant. These exclusion criteria resulted in a final sample of 27 participants for both the spatial working memory task and the math task.

We also analysed a semantic task dataset collected at the University of York, UK. 31 healthy adults were recruited from the University of York (25 females; age: mean ± SD = 21.26 ± 2.93, range: 19 – 34 years). The same exclusion criteria for functional runs were applied as above. For the feature matching task, this left 23 participants with 4 runs, 4 participants with 3 runs, and 1 participant with 2 runs. For the association task, there were 24 participants with 4 runs, 3 participants with 3 runs, and 3 participants with 2 runs. To select experimental materials, we recruited 30 native English speakers who did not participate in the main fMRI experiment as subjects (21 females; age range: 18 – 24 years). These individuals rated the color and shape similarity as well as the semantic association strength for each word pair.

### 4.2. Tasks paradigms

#### 4.2.1. Spatial working memory task

Participants were required to maintain four or eight sequentially presented locations in a 3×4 grid [28], giving rise to easy and hard spatial working memory conditions. Stimuli were presented at the center of the screen across four steps. Each of these steps lasted for 1s and highlighted one location on the grid in the easy condition, and two locations in the hard condition. This was followed by a decision phase, which showed two grids side by side (i.e., two-alternative forced choice (2AFC) paradigm). One grid contained the locations shown on the previous four steps, while the other contained one or two locations in the wrong place. Participants indicated their response via a button press and feedback was immediately provided within in 2.5s. Each run consisted of 12 experimental blocks (6 blocks per condition and 4 trials in a 32 s block) and 4 fixation blocks (each 16 s long), resulting in a total time of 448 s.

#### 4.2.2. Math task

Participants were presented with an addition expression on the screen for 1.45s and, subsequently made a 2AFC decision indicating their solution within 1s. The easy condition used single-digit numbers while the hard condition used two-digit numbers. Each trial ended with a blank screen lasting for 0.1s. Each run consisted of 12 experimental blocks (with 4 trials per block) and 4 fixation blocks, resulting in a total time of 316s.

#### 4.2.3. Semantic feature matching task

Participants were required to make a yes/no decision matching probe and target concepts (presented as words) according to a particular semantic feature (colour or shape), specified at the top of the screen during each trial. The feature prompt, probe word, and target words were presented simultaneously. Half of the trials were matching trials in which participants would be expected to identify shared features; while half of the trials were non-matching trials in which participants would not be expected to identify shared features. For example, in a colour matching trial participants would answer ‘yes’ to the word-pair DALMATIANS – COWS, due to their colour similarity, whereas they would answer ‘no’ to COAL -TOOTH as they do not share a similar colour.

This task included four runs and two conditions (two features: colour and shape), presented in a mixed design. Each run consisted of four experimental blocks (two 150s blocks per feature), resulting in a total time of 612s. In each block, 20 trials were presented in a rapid event-related design. In order to maximize the statistical power of the rapid event-related fMRI data analysis, the stimuli were presented with a temporal jitter randomized from trial to trial [31]. The inter-trial interval varied from 3 to 5 s. Each trial started with a fixation, followed by the feature, probe word, and target word presented centrally on the screen, triggering the onset of the decision-making period. The feature, probe word, and target word remained visible until the participant responded, or for a maximum of 3 s. The condition order was counterbalanced across runs and run order was counterbalanced across participants. Half of the participants pressed a button with their right index finger to indicate a matching trial and responded with their right middle finger to indicate a non-matching trial. Half of the participants pressed the opposite buttons.

#### 4.2.4. Semantic association task

Participants were asked to decide if pairs of words were semantically associated or not (i.e., yes/no decision as above) based on their own experience. Overall, there were roughly equal numbers of ‘related’ and ‘unrelated’ responses across participants. The same stimuli were used in the semantic feature matching task and semantic association task. For example, DALMATIANS and COWS are semantically related; COAL and TOOTH are not. The feature and association tasks were often separated by one week. This task included four runs, presented in a rapid event-related design. Each run consisted of 80 trials, with about half being related and half being unrelated trials. The procedure was the same as the feature matching task except only two words were presented on the screen.

### 4.3. Image acquisition

#### 4.3.1. Image acquisition for HCP dataset

MRI acquisition protocols of the HCP dataset have been previously described [30,32]. Images were acquired using a customized 3T Siemens Connectome scanner having a 100 mT/m SC72 gradient set and using a standard Siemens 32-channel radiofrequency receive head coil. Participants underwent the following scans: structural (at least one T1-weighted (T1w) MPRAGE and one 3D T2-weighted (T2w) SPACE scan at 0.7-mm isotropic resolution), rsfMRI (4 runs ×14 min and 33 s), and task fMRI (7 tasks, 46.6 min in total). Since not all participants completed all scans, we only included 339 unrelated participants from the S900 release. Whole-brain rsfMRI and task fMRI data were acquired using identical multi-band echo planar imaging (EPI) sequence parameters of 2-mm isotropic resolution with a TR = 720 ms. The dMRI data consisted of one 1.25 mm isotropic scan for each participant with 3 shell HARDI type acquisition, including b = 1000, 200, 3000 s/mm^2^, total for 270 non-collinear directions. Spin echo phase reversed images were acquired during the fMRI scanning sessions to enable accurate cross-modal registrations of the T2w and fMRI images to the T1w image in each subject and standard dual gradient echo field maps were acquired to correct T1w and T2w images for readout distortion. Additionally, the spin echo field maps acquired during the fMRI session (with matched geometry and echo spacing to the gradient echo fMRI data) were used to compute a more accurate fMRI bias field correction and to segment regions of gradient echo signal loss.

Subjects were considered for data exclusion based on the mean and mean absolute deviation of the relative root-mean-square motion across either four rsfMRI scans, resulting in four summary motion measures. If a subject exceeded 1.5 times the interquartile range (in the adverse direction) of the measurement distribution in two or more of these measures, the subject was excluded. In addition, functional runs were flagged for exclusion if more than 25% of frames exceeded 0.2 mm frame-wise displacement (FD_power). These above exclusion criteria were established before performing the analysis, (for similar implementation, see Faskowitz et al., 2020; Sporns et al., 2021). The data of 91 participants was excluded because of excessive head motion and the data of another 3 participants was excluded because their resting data did not have all the time points. In total, the data of 245 participants was analysed after exclusions.

#### 4.3.2. Image acquisition for York non-semantic dataset

MRI acquisition protocols have been described previously [14,35]. Structural and functional data were collected on a Siemens Prisma 3T MRI scanner at the York Neuroimaging Centre. The scanning protocols included a T1-weighted MPRAGE sequence with whole-brain coverage. The structural scan used: acquisition matrix of 176 × 256 × 256 and voxel size 1 × 1 × 1 mm^3^, repetition time (TR) = 2300 ms, and echo time (TE) = 2.26 ms. Functional data were acquired using an EPI sequence with an 800 flip angle and using GRAPPA with an acceleration factor of 2 in 3 x 3 x 4 mm voxels in 64-axial slices. The functional scan used: 55 3-mm-thick slices acquired in an interleaved order (with 33% distance factor), TR = 3000 ms, TE = 15 ms, FoV = 192 mm.

#### 4.3.3. Image acquisition for York semantic dataset

MRI acquisition protocols have been described previously (Wang et al., 2024a; 2024b). Whole brain structural and functional MRI data were acquired using a 3T Siemens MRI scanner utilising a 64-channel head coil, tuned to 123 MHz at York Neuroimaging Centre, University of York. The functional runs were acquired using a multi-band multi-echo (MBME) EPI sequence, each 11.45 minutes long (TR=1.5 s; TEs = 12, 24.83, 37.66 ms; 48 interleaved slices per volume with slice thickness of 3 mm (no slice gap); FoV = 24 cm (resolution matrix = 3×3×3; 80×80); 75° flip angle; 455 volumes per run; 7/8 partial Fourier encoding and GRAPPA (acceleration factor = 3, 36 ref. lines); multi-band acceleration factor = 2). Structural T1-weighted images were acquired using an MPRAGE sequence (TR = 2.3 s, TE = 2.3 s; voxel size = 1×1×1 isotropic; 176 slices; flip angle = 8°; FoV= 256 mm; interleaved slice ordering). We also collected a high-resolution T2-weighted (T2w) scan using an echo-planar imaging sequence (TR = 3.2 s, TE = 56 ms, flip angle = 120°; 176 slices, voxel size = 1×1×1 isotropic; Fov = 256 mm).

### 4.4. Image pre-processing

#### 4.4.1. Image pre-processing of HCP dataset

We used HCP’s minimal pre-processing pipelines [30]. Briefly, for each subject, structural images (T1w and T2w) were corrected for spatial distortions. FreeSurfer v5.3 was used for accurate extraction of cortical surfaces and segmentation of subcortical structures [36,37]. To align subcortical structures across subjects, structural images were registered using non-linear volume registration to the Montreal Neurological Institute (MNI152) space. Functional images (rest and task) were corrected for spatial distortions, head motion, and mapped from volume to surface space using ribbon-constrained volume to surface mapping.

Subcortical data were also projected to the set of extracted subcortical structure voxels and combined with the surface data to form the standard CIFTI grayordinate space. Data were smoothed by a 2-mm FWHM kernel in the grayordinates space that avoids mixing data across gyral banks for surface data and avoids mixing areal borders for subcortical data. Rest and task fMRI data were additionally identically cleaned for spatially specific noise using spatial ICA+FIX [38] and global structured noise using temporal ICA [39]. For accurate cross-subject registration of cortical surfaces, a multimodal surface matching (MSM) algorithm [40] was used to optimize the alignment of cortical areas based on features from different modalities. MSMSulc (“sulc”: cortical folds average convexity) was used to initialize MSMAll, which then utilized myelin, resting state network, and rfMRI visuotopic maps. Myelin maps were computed using the ratio of T1w/T2w images [38]. The HCP’s minimally preprocessed data include cortical thickness maps (generated based on the standardized FreeSurfer pipeline with combined T1-/T2-reconstruction). For this study, the standard-resolution cortical thickness maps (32k mesh) were used.

#### 4.4.2. Image pre-processing of York non-semantic and semantic dataset

The York datasets were preprocessed using fMRIPrep 20.2.1 [[41], RRID:SCR_016216], with detailed methods previously described in [6,23]. In brief, anatomical preprocessing involved intensity non-uniformity correction, skull stripping, segmentation, and surface reconstruction. Spatial normalization to MNI152 templates was performed through nonlinear registration. For each BOLD run, functional data preprocessing followed standard fMRIPrep procedures. This included generating a reference volume, applying fieldmap correction, and co-registering the BOLD reference to anatomical space. Motion correction and slice-time correction were performed. Resampling was done in the surface space - fsaverage, and final data were output in CIFTI grayordinate space. Post-processing of fMRIPrep outputs was performed using eXtensible Connectivity Pipeline (XCP) [42]. Volumes with framewise displacement greater than 0.3 mm were excluded before nuisance regression. A total of 36 nuisance regressors, including motion parameters, global signal, white matter, and CSF signals, were regressed. Residual time-series were band-pass filtered (0.01-0.08 Hz) and smoothed with a 6.0 mm FWHM Gaussian kernel. Detailed information is provided in Supplementary Materials and Methods, Section 1.1.

### 4.5. Task fMRI analysis

#### 4.5.1. Individual-specific parcellation

To account for anatomical and functional variability, we used a multi-session hierarchical Bayesian model to estimate individual-specific parcellations, following the methods in [6,19]. This approach defines 400 individualized parcels across 17 networks per participant and the individual-specific parcellation showed greater homogeneity than the parcellation using group atlas [6,19]. All subsequent analyses were based on parcel-specific time-series. Detailed information is provided in Supplementary Materials and Methods, Section 1.2.1.

#### 4.5.2. Constructing fMRI FC matrices

To investigate how FC varies across tasks, we computed task-based and resting state FC matrices. We opted not to use the traditional psychophysiological interaction method for measuring task-state functional connectivity due to its potential to inflate activation-induced task-state functional connectivity, which may identify regions that are active rather than interacting during the task [43]. Since task activations can spuriously inflate task-based functional connectivity estimates, it is necessary to correct for task-timing confounds by removing the first-order effect of task-evoked activations (i.e., mean evoked task-related activity, likely active during the task) prior to estimating task-state functional connectivity (likely interacting during the task) [43]. Specifically, we fitted the task timing for each task using a finite impulse response model, a method that has been shown to reduce both false positives and false negatives in FC estimation [44–46]. In semantic tasks, approximately 5 timepoints were modeled for each trial. For the non-semantic task, roughly 2.5 timepoints were modeled for each trial.

Following task regression, we demeaned the residual time series for each parcel and quantified the FC using Pearson correlation for each participant, task and run. The Pearson correlation coefficients might be inflated due to the temporal autocorrelation in task fMRI time series data [47]. To address this, we corrected the Pearson correlation using a correction approach, xDF, which accounts for both autocorrelation within each time series as well as instantaneous and lagged cross-correlations between the time series [48]. This method provides an effective degrees of freedom estimator that addresses cross-correlations, thereby preventing inflation of Pearson correlation coefficients. Our goal was not to remove temporal autocorrelation, but to enhance the precision and reliability of correlation estimates, reducing false functional connectivity between regions. We calculated xDF-adjusted z-scored correlation coefficients to assess the interregional relationships in BOLD time series, resulting in a 400 × 400 functional connectivity matrix for each participant, task, and run. Finally, we averaged these functional connectivity estimates within networks, and between pairs of networks, to construct a network-by-network functional connectivity matrix. The same method was used to calculate the resting-state functional connectivity of the HCP dataset and construct a corresponding network-by-network functional connectivity matrix, except without the task regression step.

#### 4.5.3. Feature extraction of the time-series data

To investigate how the FS patterns vary across tasks, we calculated task-based and resting state FS using the extracted features. To extract the features of the time-series required for this analysis, we used the time-series analysis toolbox (hctsa) [8,49]. With this tool, we transformed each time-series in the dataset into a set of over 7,700 features, which include, but are not limited to, distributional properties, entropy and variability, autocorrelation, time-delay embeddings, and nonlinear properties of a given time-series (Fig. 1) [8,9]. We extracted features from the parcellated fMRI time-series of each participant, each task, and each run separately. After the feature extraction procedure, we removed the outputs of the operations that produced errors and normalized the remaining features (about 6900 features) across parcels using an outlier-robust sigmoidal transform. The resulting normalized feature matrix (400 parcels × ∼7000 features × 4 runs) was used to predict networks labels of parcels to identify the DAN subnetwork with varying interaction patterns across tasks (Method 4.5.4) and to construct FS matrix for further analysis (Method 4.5.4).

#### 4.5.4. Constructing fMRI FS matrices

To investigate how FS patterns vary across tasks, we calculated task-based and resting state FS matrices. Specifically, we calculated Pearson correlation coefficients of the extracted features, which represented the pairwise FS between all possible combinations of brain parcels (Fig. 1). This resulted in a 400 by 400 FS matrix for each run each task for each participant. Finally, to construct a network-by-network FS matrix, we averaged the estimates of FS within networks and between pairs of networks for further analysis.

To investigate whether functionally connected regions display similar features, we calculated Pearson correlations coefficients for FS within and between networks defined by resting state FC. Then we tested whether the correlations for FS within networks were greater than the correlations between networks.

Additionally, we estimated the similarity between FC and FS by calculating their correlations at rest and for each task. To control for multiple comparison, we FWE-corrected the p values using permutation-based maximum r values. Specifically, we created a null distribution using permutation for each task and chose the maximum values among all the tasks. We then compared the observed correlation value with the null distribution to examine whether the correlation between FC and FS was significantly greater than that expected from null distribution.

#### 4.5.5. Dimension reduction analysis of FC and FS

To examine whether FC and FS captured similar principal components, we performed dimension reduction analysis on both resting state FC matrix and FS matrix derived from the HCP dataset. For the FC matrix, we first calculated resting state FC for each run of each participant, as detailed in Method section 2.5.2. Subsequently, these individual connectivity matrices were then averaged to calculate a group-level connectivity matrix. We extracted ten group-level gradients from the group-level connectivity matrix (dimension reduction technique = diffusion embedding, kernel = None, sparsity = 0.9), in line with previous studies [35,50] using the Brainspace Toolbox [51]. This analysis resulted in ten group-level gradients explaining maximal whole-brain connectivity variance in descending order. A parallel analysis was performed for the FS matrix, employing identical procedures except for the input, which consisted of the FS matrix as delineated in Method Section 2.5.4. This analysis resulted in ten group-level gradients explaining maximal whole-brain FS variance in descending order. We retained the components explaining the most variance by looking at the eigenvalues of each component in the scree plots shown in Fig 3.

Finally, we examined the similarities between the first three components captured by FC and FS, respectively via calculating the Pearson correlations between corresponding components. Due to the spatial autocorrelation present in each principal component, we created a null distribution using spin permutation implemented in BrainSMASH [52]. This approach simulates brain maps, constrained by empirical data, that preserve the spatial autocorrelation of cortical parcellated brain maps. We then compared the observed correlation value with the null distribution for left hemisphere to examine whether the correlation between two components of parcels in the left hemisphere was significantly greater than that expected from spatial autocorrelation alone. Similarly, we examined the correlation between two components of parcels in the right hemisphere using the same methods. This analysis was performed for the two hemispheres separately because the geodesic distance between parcels was used to generate the spatial-autocorrelation-preserving surrogate maps when creating the null distribution. It was only possible to measure within-hemisphere geodesic distance between parcels because the left and right hemisphere surface maps were not on the same mesh.

#### 4.5.6. Timescale analysis

The first component captured by FS matrix was similar to the timescale gradient map reported before [7], as determined by visual comparison. To further understand this component, we calculated its correlation with the intrinsic timescale gradient map, which represents the temporal duration of ongoing inputs that the brain can process. The intrinsic timescale for each parcel was characterized by the decay of the temporal autocorrelation function, as the time taken for the autocorrelation function to reach a threshold of r = 0.5 (i.e., half of the full width at half maximum), consistent with prior studies [7,53]. A higher value of the intrinsic timescale indicates longer ongoing inputs that the parcel can process. Given the spatial autocorrelation present in this component and the timescale gradient map, we created a null distribution using spin permutation implemented in BrainSMASH [52]. We subsequently compared the observed correlation value with the null distribution for each hemisphere to determine whether the real correlations were significantly greater than that expected by spatial autocorrelation alone.

#### 4.5.7. Calculating the correlations across tasks in FC and in FS

To test whether FS was more sensitive to task modulation than FC, we calculated correlations across tasks for both measures and compared their sensitivities by assessing if FS showed weaker correlations than FC. We used two methods: (i) we first calculated Pearson correlations (r1) for FC across tasks for each possible pair of tasks at the group level (for instance, between the spatial working memory and math tasks) using the task mean FC matrices. Then, we repeated the process for FS (r2), using the task mean FS matrices. Finally, we then compared the resulting correlation values (r1 versus r2). (ii) We calculated the Pearson correlation coefficients between the FC matrix of the spatial working memory task and the FC matrix of math task at the individual level for each participant given that each participant completed both spatial working memory task and math task. Then we repeated the process for FS, using the FS matrix of each participant. Finally, we converted the r values to z values using Fisher transformation and compared the z values of the FC with the ones of the FS by conducting paired t-tests. The same procedures were applied to the two semantic tasks (semantic feature matching versus association).

#### 4.5.8. Classification analysis – decoding task labels using FS matrix

To determine whether feature similarity captures task information, we conducted classification analyses to predict task labels (spatial working memory versus math) using FS matrices of the non-semantic tasks of all the participants, respectively. We employed scikit-learn’s [54] linear support vector machine classifier (SVC) with 5-fold cross-validation to avoid overfitting and ensure reliable performance. To assess statistical significance, we performed permutation tests by shuffling task labels 1,000 times, creating a null distribution for comparison [55]. The same approach was used for semantic tasks (semantic feature matching versus. semantic association) using feature similarity matrices of semantic tasks.

#### 4.5.9. Comparing FC difference and FS difference between networks across tasks

To investigate whether FS captured varying interaction patterns across tasks that could not be captured by FC, we first calculated the average FC between DAN-A and Visual network and between DAN-A and DMN, across all runs per participant per task. Subsequently, we calculated the relative FC difference by subtracting the FC between DAN-A and DMN from the FC between DAN-A and Visual network. Finally, we conducted paired-t tests for each task to examine the significance of the FC difference between network pairs. We further investigated whether the task influenced the FC difference between network pairs using the maximum/minimum permutation test. We calculated the mean FC difference between DAN-A and Visual network versus DAN-A and DMN for each task and calculated the mean FC difference between each task pair. To access statistical significance, we permutated the task label 10000 times and then calculated the mean FC difference between these two tasks to build a null distribution for each task pair. To control the family-wise error (FWE) rate (p = 0.05, FWE-corrected) given the inclusion of multiple task pairs, we utilized the permutation-based maximum mean FC difference and minimum mean FC difference values in the null distribution for each task pair. To evaluate significance, if the observed mean difference value was positive, we counted the percentage of times that mean difference values in the maximum null distribution were greater than the observed ‘true’ mean difference values. Conversely, if the observed mean difference value was negative, we counted the percentage of times of mean difference values in the minimum null distribution were less than the observed ‘true’ mean difference values.

We then examined whether task influences the FS differences between targeted network pairs (DAN-A-Visual versus DAN-A-DMN) using the same procedures but applied to FS. Our results showed that FS is more sensitive to task modulation than FC in the DAN-A-Visual and DAN-A-DMN comparison. To assess whether this holds for other networks, we repeated the analysis, comparing FPCN-A-Visual versus FPCN-A-DMN.

#### 4.5.10. Classification analysis – decoding network labels of parcels

Given heterogeneity of DAN, we used a data-driven approach to identify the subnetwork most likely to vary in interaction patterns during tasks using the extracted features. Specifically, we conducted a classification analysis on a normalized feature matrix (400 parcels by about 7000 features by 4 runs) to decode parcel network labels. After confirming accurate classification, we examined the confusion matrix to identify classification errors (Detailed information is provided in Supplementary Materials and Methods, Section 1.2.3). This allowed us to explore network similarity, as functionally similar networks are more likely to be misclassified.

#### 4.5.11. Constructing fMRI statistics of pairwise interaction matrices

Many techniques have been developed to measure pairwise interactions in complex systems, such as Pearson correlation in fMRI and mutual information in signal processing. These methods, which range from contemporaneous correlation coefficients to causal inference approaches, are based on distinct quantitative theories and define interactions differently. As a result, each method captures different information and has varying sensitivity. However, since most methods focus on a single aspect, their sensitivity may still be weaker than FS, which considers multiple dimensions of information. To test this, we calculated many SPIs using the Python Toolkit for Statistics for Pairwise Interactions (pyspi; v0.4.1; [1]. The parcel-based task fMRI time series were z-scored along the time dimension before the calculation.

Different systems involve distinct interactions, causing SPIs to perform variably across datasets (e.g., smartwatch activity, EEG, fMRI), with some methods excelling only in specific datasets [1]. Therefore, it’s essential to select SPIs that capture the relevant interactions for a given dataset. From the original list of SPIs [1], we initially selected all 67 SPIs that significantly classified states in the fMRI film dataset and then refined this set to 49 SPIs with reasonable computational requirements (<5 hours per run per task per subject). This approach ensured the selected SPIs were both relevant and practical for our analysis.

The 49 SPIs used in this study were derived from 20 interaction measures across six categories: (1) Basic Statistics: This category includes measures such as covariance, which quantifies the linear relationship between two variables, and precision, which captures pairwise associations while controlling for the effects of other variables in the dataset. (2) Distance-Based Similarity: These measures quantify statistical similarity or independence using pairwise distances between bivariate observations. Examples include Euclidean distance and correlation distance, which evaluate how similar or dissimilar two variables are in their distributions. (3) Information-Theoretic Measures: Metrics in this category include mutual information, which quantifies the total dependency between two variables, and joint entropy, which measures the combined uncertainty or information content of two variables. (4) Causal Inference: Methods in this category are designed to infer directed relationships. For example, regression error-based causal inference determines the causal direction by analysing which variable’s residuals are more independent of the predictor, indicating potential causal influence. (5) Spectral Measures: These measures are computed in the frequency or time-frequency domain. Examples include spectral coherence magnitude, which quantifies the alignment of frequency components in phase and amplitude, and phase locking value, which measures the consistency of phase differences between two signals over time. (6) Miscellaneous Methods: This category includes techniques such as linear model fits, which estimate relationships through regression, and other statistical tools that quantify pairwise interactions but do not fit into the above categories. See SM 1.3.3 for a full list of SPIs used in this study.

We calculated the pairwise statistics for the selected SPIs across all subjects and compared the interaction matrices across tasks, focusing on key networks (DAN-A, Visual, and DMN). We conducted paired t-tests for each task to assess differences between network pairs and conducted FDR correction to control for multiple comparisons. The same methods were used when comparing FC and FS, with the only difference being the input matrices derived from SPIs.

### 4.6. Data and code Accessibility

The HCP dataset is publicly available at https://www.humanconnectome.org/. Raw data collected at the University of York cannot be shared at this time due to consent limitations under UK GDPR. Researchers interested in accessing this data should contact the Chair of the Research Ethics Committee at the York Neuroimaging Centre. Preprocessed data from the University of York are available on the Open Science Framework at https://osf.io/t2p7j/, and analysis code is accessible at https://github.com/Xiuyi-Wang/Feature_similarity_Project.

## Abbreviations

DAN: dorsal attention network;
DMN: default mode network;
FC: functional connectivity,
FPCN: fronto-parietal control network;
FS: feature similarity;
SPI: statistics of pairwise interaction

## Acknowledgements

We are grateful to Pradeepa Ruwan and Antonia De Freitas for piloting the experiment. We would like to thank Ben D Fulcher for providing codes and a guide to help us extract the features using hctsa. X. W. discloses support for the research of this work from National Natural Science Foundation of China (Grant Number. 32300881). Y. D. discloses support for the publication of this work from the STI 2030—Major Projects (Grant Number. 2021ZD0201500), the National Natural Science Foundation of China (Grant No. 31822024), and Scientific Foundation of Institute of Psychology, Chinese Academy of Sciences (Grant Number. E2CX3625CX). E. J. discloses support for the research of this work from a European Research Council Consolidator grant (Project ID: 771863 - FLEXSEM). N. L. discloses support for the research of this work from National Natural Science Foundation of China (Grant Number. 32471111).

