## Supplemental text and figure for "Feature similarity: a sensitive method to capture the functional interaction of brain regions and networks to support flexible behavior"

Short title: Feature Similarity: Capturing Brain Network Interactions for Flexible Behavior

Xiuyi Wang[1,2,3]*, Baihan Lyu[1,2, 3], Katya Krieger-Redwood[4], Nicholas E. Souter[4], Golia Shafiei[5], Nan Lin[1,2,3], Jonathan Smallwood[6], Elizabeth Jefferies[4]*, Yi Du[1,2,3,7]*

1. State Key Laboratory of Cognitive Science and Mental Health, Institute of Psychology, Chinese Academy of Sciences, Beijing 100101, China

2. Institute of Psychology, Chinese Academy of Sciences, Beijing, 100101, China.

3. Department of Psychology, University of Chinese Academy of Sciences, Beijing 100049, China.

4. Department of Psychology, University of York, Heslington, York, YO10 5DD, United Kingdom.

5. Department of Psychiatry, Perelman School of Medicine, University of Pennsylvania, Philadelphia, PA 19104, USA.

6. Department of Psychology, Queens University, Kingston, Ontario, Canada.

7. Chinese Institute for Brain Research, Beijing 102206, China.

### Supplemental materials

### 1. Materials and Methods

#### 1.1. Image pre-processing of York dataset

##### 1.1.1. Anatomical data preprocessing

The T1w image was corrected for intensity non-uniformity (INU) with N4BiasFieldCorrection [1], distributed with ANTs 2.3.3 [[2], RRID:SCR_004757], and used as T1w-reference throughout the workflow. The T1w-reference was then skull-stripped with a Nipype implementation of the antsBrainExtraction.sh workflow (from ANTs), using OASIS30ANTs as target template. Brain tissue segmentation of cerebrospinal fluid (CSF), white-matter (WM) and gray-matter (GM) was performed on the brain-extracted T1w using fast FSL 5.0.9[[3], RRID:SCR_002823]. Brain surfaces were reconstructed using recon-all from FreeSurfer 6.0.1 [(Dale et al. 1999), RRID:SCR_001847], and the brain mask estimated previously was refined with a custom variation of the method to reconcile ANTs-derived and FreeSurfer-derived segmentations of the cortical gray-matter of Mindboggle [[5], RRID:SCR_002438]. Volume-based spatial normalization to two standard spaces (MNI152NLin2009cAsym, MNI152NLin6Asym) was performed through nonlinear registration with antsRegistration (ANTs 2.3.3), using brain-extracted versions of both T1w reference and the T1w template. The following templates were selected for spatial normalization: ICBM 152 Nonlinear Asymmetrical template version 2009c [[6], RRID:SCR_008796; TemplateFlow ID: MNI152NLin2009cAsym], FSL's MNI ICBM 152 non-linear 6th Generation Asymmetric Average

Brain Stereotaxic Registration Model [[7], RRID:SCR_002823; TemplateFlow ID: MNI152NLin6Asym].

1.1.2. Functional data preprocessing

For each of the BOLD runs per subject, the following preprocessing was performed. First, a reference volume and its skull-stripped version were generated using a custom methodology of fMRIPrep. A B0-nonuniformity map (or fieldmap) was estimated based on a phase-difference map calculated with a dual-echo GRE (gradient-recall echo) sequence, processed with a custom workflow of SDCFlows inspired by the epidewarp.fsl script and further improvements in HCP Pipelines [8]. The fieldmap was then co-registered to the target EPI reference run and converted to a displacements field map (amenable to registration tools such as ANTs) with FSL's fugue and other SDCflows tools. Based on the estimated susceptibility distortion, a corrected EPI reference was calculated for a more accurate co-registration with the anatomical reference. The BOLD reference was then co-registered to the T1w reference using bbregister (FreeSurfer) which implements boundary-based registration [9]. Co-registration was configured with six degrees of freedom. Head-motion parameters with respect to the BOLD reference (transformation matrices, and six corresponding rotation and translation parameters) are estimated before any spatiotemporal filtering using mcflirt (FSL 5.0.9, Jenkinson et al., (2002). BOLD runs were slice-time corrected using 3dTshift from AFNI 20160207 [(27), RRID:SCR_005927]. The BOLD time-series were resampled onto the following surfaces (FreeSurfer reconstruction nomenclature): fsaverage. Grayordinates files [8] containing 91k samples were also generated using the highest-resolution fsaverage as intermediate standardized surface space. Several confounding time-series were calculated based on the preprocessed BOLD: framewise displacement (FD), DVARS and three region-wise global signals. FD was computed using two formulations following Power (absolute sum of relative motion; Power et al., 2014, relative root mean square displacement between affines; Jenkinson et al., 2002). FD and DVARS were calculated for each functional run, both using their implementations in Nipype (following the definitions by Power et al. (2014). The three global signals were extracted within the CSF, the WM, and the whole-brain masks. Additionally, a set of physiological regressors were extracted to allow for component-based noise correction (CompCor, Behzadi et al., (2007), principal components were estimated after high-pass filtering the preprocessed BOLD time-series (using a discrete cosine filter with 128s cut-off) for the two CompCor variants: temporal (tCompCor) and anatomical (aCompCor). tCompCor components were then calculated from the top 2% variable voxels within the brain mask. For aCompCor, three probabilistic masks (CSF, WM and combined CSF+WM) are generated in anatomical space. The implementation differs from that of Behzadi et al. (2007) in that instead of eroding the masks by 2 pixels in BOLD space, the aCompCor masks are subtracted from a mask of pixels that likely contain a volume fraction of GM. This mask is obtained by dilating a GM mask extracted from the FreeSurfer’s aseg segmentation, and it ensures components are not extracted from voxels containing a minimal fraction of GM. Finally, these masks are resampled into BOLD space and binarized by thresholding at 0.99 (as in the original implementation). Components are also calculated separately within the WM and CSF masks. For each CompCor decomposition, the k components with the largest singular values are retained, such that the retained components’ time-series are sufficient to explain 50 percent of variance across the nuisance mask (CSF, WM, combined, or temporal). The remaining components are dropped from consideration. The head-motion estimates calculated in the correction step were also placed within the corresponding confounds file. The confound time-series derived from head motion estimates and global signals were expanded with the inclusion of temporal derivatives and quadratic terms for each [14]. Frames that exceeded a threshold of 0.5 mm FD or 1.5 standardized DVARS were annotated as motion outliers. All resamplings can be performed with a single interpolation step by composing all the pertinent transformations (i.e., head-motion transform matrices, susceptibility distortion correction when available, and co-registrations to anatomical and output spaces). Gridded (volumetric) resamplings were performed using antsApplyTransforms (ANTs), configured with Lanczos interpolation to minimize the smoothing effects of other kernels [15]. Non-gridded (surface) resamplings were performed using mri_vol2surf (FreeSurfer). Many internal operations of fMRIPrep use Nilearn 0.6.2 [Abraham et al., (2014) RRID:SCR_001362], mostly within the functional processing workflow. To resulting data were in CIFTI 64k-vertex grayordinate space. The left hemisphere has 29696 vertices and right hemisphere has 29716 vertices in total after removing the medial wall.

The post-processing of the outputs of fMRIPrep version 20.2.1 [18] was performed using the eXtensible Connectivity Pipeline (XCP) [14,19]. XCP was built with Nipype 1.7.0 [20]. For each CIFTI run found per subject, the following post-processing was performed: before nuisance regression and filtering any volumes with framewise-displacement greater than 0.3 mm [12,14] were flagged as outlier and excluded from nuisance regression. In total, 36 nuisance regressors were selected from the nuisance confound matrices of fMRIPrep output. These nuisance regressors included six motion parameters, global signal, the mean white matter, the mean CSF signal with their temporal derivatives, and the quadratic expansion of six motion parameters, tissues signals and their temporal derivatives [14,19,21]. These nuisance regressors were regressed from the BOLD data using linear regression - as implemented in Scikit-Learn 0.24.2 [22]. Residual time-series from this regression were then band-pass filtered to retain signals within the 0.01-0.08 Hz frequency band. The processed BOLD was smoothed using Connectome Workbench with a gaussian kernel size of 6.0 mm (FWHM). Processed functional time-series were extracted from residual BOLD using Connectome Workbench [8]: for the Glasser atlas [23]. Many internal operations of XCP use Nibabel [16], numpy [24], and scipy [24].

### 1.2. Task fMRI analysis

#### 1.2.1. Individual-specific parcellation

To account for anatomical and functional variability, we used a multi-session hierarchical Bayesian model (MS-HBM) to estimate individual-specific parcellations, following the methods in [25,26]. Pseudo-resting state time-series were generated by regressing out task activations from fMRI data. This approach defines 400 individualized parcels across 17 networks per participant, incorporating both local gradients and spatial contiguity priors to improve homogeneity compared to group-level atlases. All subsequent analyses were based on parcel-specific time-series.

Considering the anatomical and functional variability across individuals, we estimated individual-specific areal-level parcellation using a multi-session hierarchical Bayesian model

(MS-HBM), see Kong et al. (2021, 2019) for the details of the method; see SM 1.2.1 and (Wang et al. (2024) for the details of the application of this method to these three datasets and the greater homogeneity of individual-specific parcellation than the parcellation using group atlas. To estimate individual-specific parcellation, we acquired “pseudo-resting state” time-series in which the task activation model was regressed from feature matching, semantic association, spatial working memory, and math fMRI data [28] using xcp_d (https://github.com/PennLINC/xcp_d). The task activation model and nuisance matrix were regressed out using AFNI’s3dTproject (for similar implementation, see Cui et al. (2020)).

Using a group atlas, this method calculates inter-subject resting state FC variability, intra-subject resting state FC variability, and finally parcellates for each single subject based on this prior information. MS-HBM requires FC profiles of multiple sessions as input; here, we included two separate sessions, with each including four fMRI runs for each York dataset (we included another feature matching task which has 4 runs in York non-semantic dataset to generate individual-specific parcellation). As in Kong et al. (2021, 2019), we used MS-HBM to define 400 individualized parcels belonging to 17 discrete individualized networks for each participant. Specifically, we calculated all participants’ connectivity profiles, created the group parcellation using the average connectivity profile of all participants, estimated the inter-subject and intra-subject connectivity variability, and finally calculated each participant’s individualized parcellation. This parcellation imposed the Markov random filed (MRF) spatial prior. We used a well-known areal-level parcellation approach, i.e., the local gradient approach (gMS-HBM), which detects local abrupt changes (i.e., gradients) in resting state FC across the cortex [30]. A previous study [31] has suggested combining local gradient [30,32] and global clustering (Yeo et al., 2011) approaches for estimating areal-level parcellations. Therefore, we complemented the spatial contiguity prior in contiguous MS-HBM (cMS-HBM) with a prior based on local gradients in resting state FC, which encouraged adjacent brain locations with gentle changes in FC to be grouped into the same parcel. We used parameters (i.e., beta value = 50, w = 30 and c = 30) optimized using our own dataset. The same parameters were also used in Kong et al. (2021). Vertices were parcellated into 400 cortical regions (200 per hemisphere). To parcellate each of these parcels, we calculated the average time-series of enclosed vertices to get better signal noise ratio (SNR) using the Connectome Workbench software. All the following analysis used this parcel-based time-series. The same method and parameters were used to generate the individual-specific parcellation for the participants in the HCP dataset using the resting state time-series except that the task regression was not performed.

##### 1.2.1.1 Homogeneity of parcels

To evaluate whether the functional parcellation was successful, we examined parcel homogeneity [25,26,32]. This was calculated as the average Pearson's correlation coefficient between fMRI time courses of all pairs of vertices within each parcel, adjusted for parcel size and summed across parcels [25,26,31]. Higher homogeneity reflects that vertices within the same parcel share more similar time courses indicating higher parcellation quality. To summarize parcel homogeneity, we averaged the homogeneity value across parcels. We calculated the parcel homogeneity for each run of each task for a given participant using their individual-specific parcellation and then averaged them across runs for each participant for each task. We also calculated the parcel homogeneity using the canonical Yeo 17-network group atlas. Using the resting state HCP data, Kong et al. (2021) demonstrated that homogeneity within MS-HBM-based individualized parcels was greater than in the canonical Yeo 17-network group atlas, which does not consider variation in functional neuroanatomy. A similar pattern, (i.e., the individual-specific parcellation showed greater homogeneity than the parcellation using group atlas), was observed for the York non-semantic and semantic datasets[27].

##### 1.2.2. Classification analysis – decoding the task labels

To explore whether FS captured meaningful task information, we conducted classification analyses to access the accuracy with which a classifier could predict the task labels of non-semantic tasks (i.e., spatial working memory task versus math task) using FS matrices of the non-semantic tasks of all the participants. We performed the two-class classification (spatial working memory task versus math task) using scikit-learn (Pedregosa et al., 2011), which is a general machine learning library written in Python. We trained linear support vector machine classifiers (sklearn.svm.SVC) to identify the hyperplane that maximally separates the samples belonging to different classes. We utilized 5-fold cross-validation to minimize the risk of overfitting and to obtain reliable performance estimates. To determine whether the classification accuracy was significantly greater than chance level, we performed the permutation-based multi-class classification analysis in which we randomly shuffled the task labels 1000 times within all participants. This established an empirical distribution of classification accuracy scores under the null hypothesis where there is no association between FS matrices and task labels [34]. We employed similar procedures to investigate the accuracy with which a classifier could predict the task labels of semantic tasks (i.e., semantic feature matching versus semantic association task) using FS matrices of semantic tasks. We also performed similar classification analysis using FC matrices.

#### 1.2.3. Classification analysis – decoding the network labels of parcels

To identify the subnetwork of DAN that is more likely to vary interaction patterns during tasks, we employed a data-driven approach to characterize the network similarity of DAN. Specifically, we conducted a classification analysis using a normalized feature matrix. Previous studies have established that Visual network, DAN, and FPCN consist of a multiple demand network that supports cognitive function across tasks [35,36] and these networks exhibit changes in their interaction patterns in response to task demands [27,37]. Despite the heterogeneous nature of both DAN and FPCN [27,38], it is established that FPCN-A subnetwork displays positive FC with DAN and visual networks, while showing negative FC with the DMN [27,38,39]. However, it remains unclear which subnetwork of DAN exhibits a stronger relationship with Visual network. To address this question, we conducted a classification analysis to decode the network labels of parcels. Specifically, we examined whether DAN-A exhibited a stronger relationship with Visual network than DAN-B by testing whether compared with DAN-B, DAN-A was more likely to be misclassified as Visual network. We also examined whether DAN-A was most likely to be misclassified as Visual network but DAN-B was not.

1.2.3.1. Balanced accuracy of classification analysis

To investigate whether we could classify the network labels of each parcel using the extracted features, we performed a classification analysis. The detailed information was reported in [27] and pasted here. Specifically, we investigate how accurately a classifier can learn a mapping from time-series features of parcels to network labels of these parcels. To accomplish this, we combined the normalized features from all runs of each task for each participant and performed multi-class classification (17 networks labels) using scikit-learn, a general machine learning library written in Python [22].

For multi-class classification, we trained linear support vector machine classifiers (sklearn.svm.SVC) to find the hyperplane that maximally separates the samples belonging to different classes. As the number of parcels in networks varies (e.g., both FPCN-A and FPCN-B have 25 parcels, while FPCN-C has 23 parcels), we reported balanced classification accuracy to account for the imbalance of observations across networks [40,41]. Specifically, balanced accuracy was calculated as the arithmetic mean of sensitivity (i.e., true positive rate, which measures the proportion of correctly predicted positives), and specificity, (i.e., true negative rate, which measures the proportion of correctly identified negatives). To prevent overfitting and optimistic performance estimates, we performed 5-fold cross-validation employing a within-subject cross-validation approach. We merged data from all runs per task per participant, creating datasets with 1600 samples (400 parcels × 4 runs) for semantic tasks and 800 samples (400 parcels × 2 runs) for non-semantic tasks. This dataset underwent a 5-fold cross-validation, with data from the same and different runs used in training and testing phases. To access whether the classification accuracy was significantly above chance level for each participant and each task, we performed permutation-based multi-class classification analysis by randomly shuffling network labels 1000 times within all runs for each participant for each task. This established an empirical distribution of classification accuracy scores under the null hypothesis where there is no association between features and network labels [34].

Given the high dimensionality of the data used for classification (approximately 7000 features and 1600 samples for semantic tasks; and 800 samples for non-semantic tasks), machine learning with many more features than samples are challenging. We addressed the challenge posed by the curse of dimensionality, which describes the increase in noise and redundancy with the explosive nature of increasing data dimensions [42]. To explore the curse of dimensionality, we performed classification using all the features and subsets of features, ranging from 500 to 7000 features, with increments of 500 features. Subsets of features were the top features that make major contributions in the classification when including all the features. We observed minimal accuracy cost with fewer features and no curse of dimensionality. Specifically, the accuracy increased with the number of features, peaking at 4000 features, beyond which a slight decrease was noted but remained above chance level. Consequently, we used 4000 features for the classification accuracy and confusion matrix presented in the main text.

1.2.3.2. Confusion matrix for classification analysis

The classification accuracy allowed us to evaluate whether we could correctly classify the network labels of parcels. However, classification accuracy alone can obscure the detail needed to diagnose the performance of our model. For example, for multi-class classification, high classification accuracy may be observed because all classes are being predicted equally well or because one or two classes are being neglected by the model. Therefore, to better understand the performance of the classification model, we analyzed the confusion matrix, which summarizes the prediction results. In the confusion matrix, a row represents an instance of the actual class (i.e., an actual network), whereas a column represents an instance of the predicted class (i.e., the predicted network). The diagonal elements represent the number of points for which the predicted label is equal to the true label, while off-diagonal elements are those that are mislabeled by the classifier. Higher diagonal values indicate more correct predictions. The confusion matrix reveals not only the errors made by the classifier but also the types of errors made, showing the ways in which our classification model is confused when it makes predictions. Analyzing the classification output allows us to explore the network similarity because networks with more similar functions are more likely to be incorrectly classified as each other.

To account for the imbalance of observations across the networks, we normalized the confusion matrices by the total number of elements in each class. Here we focus on DAN and reported the normalized confusion matrix for each subnetwork of DAN. We examined whether the percentage of predicted networks were different between networks by conducting paired t-tests and performed FDR correction at $p = 0.05$ to control for multiple comparisons. The normalized confusion matrix of FPCN has been reported in [27], with FPCN-A showed greater feature similarity with DAN-A and visual network.

##### 1.3.3. Statistics of pairwise interactions (SPIs)

From the original list of SPIs [1], we initially selected all 67 SPIs that significantly classified states in the fMRI film dataset and then refined this set to 49 SPIs with reasonable computational requirements (<5 hours per run per task per subject). The full list of SPIs can be found in Table 1. See Cliff et al. (2023) [43] for the explanation of each SPI.

**Table 1.** The list of 49 SPIs included in this study.

| Index | Statistics | Description | Category |
|---|---|---|---|
| 1 | cov_EllipticEnvelope | Covariance | Basic statistics |
| 2 | cov_EmpiricalCovariance | Covariance | Basic statistics |
| 3 | cov_GraphicalLassoCV | Covariance | Basic statistics |
| 4 | cov_LedoitWolf | Covariance | Basic statistics |
| 5 | cov_MinCovDet | Covariance | Basic statistics |
| 6 | cov_OAS | Covariance | Basic statistics |
| 7 | cov_ShrunkCovariance | Covariance | Basic statistics |
| 8 | prec_EllipticEnvelope | Precision | Basic statistics |
| 9 | prec_EmpiricalCovariance | Precision | Basic statistics |
| 10 | prec_GraphicalLassoCV | Precision | Basic statistics |
| 11 | prec_LedoitWolf | Precision | Basic statistics |
| 12 | prec_MinCovDet | Precision | Basic statistics |
| 13 | prec_OAS | Precision | Basic statistics |
| 14 | prec_ShrunkCovariance | Precision | Basic statistics |
| 15 | spearmanr | Spearman's rank-correlation coefficient | Basic statistics |
| 16 | kendalltau | Kendall's rank-correlation coefficient | Basic statistics |
| 17 | reci | Regression error-based causal | Causal inference |

| | | | |
|---|---|---|---|
| | | inference | |
| 18 | bary_euclidean_max | Barycenter | Distance similarity |
| 19 | dcorr | Cross distance correlation | Distance similarity |
| 20 | dcorr_biased | Cross distance correlation | Distance similarity |
| 21 | dtw | Dynamic time warping | Distance similarity |
| 22 | dtw_constraint-itakura | Dynamic time warping | Distance similarity |
| 23 | dtw_constraint-sakoe-chiba | Dynamic time warping | Distance similarity |
| 24 | hsic | Hilbert-Schmidt Independence Criterion | Distance similarity |
| 25 | hsic_biased | Hilbert-Schmidt Independence Criterion | Distance similarity |
| 26 | lcss | Longest common subsequence | Distance similarity |
| 27 | lcss_constraint-sakoe-chiba | Longest common subsequence | Distance similarity |
| 28 | softdtw | Soft dynamic time warping | Distance similarity |
| 29 | softdtw_constraint-itakura | Soft dynamic time warping | Distance similarity |
| 30 | softdtw_constraint-sakoe-chiba | Soft dynamic time warping | Distance similarity |
| 31 | cce_kernel_W-0.5 | Causally conditioned entropy | Information theory |
| 32 | cce_gaussian | Causally conditioned entropy | Information theory |
| 33 | ce_gaussian | Conditional entropy | Information theory |
| 34 | ce_kernel_W-0.5 | Conditional entropy | Information theory |
| 35 | je_gaussian | Joint entropy | Information theory |
| 36 | mi_gaussian | Mutual information | Information theory |
| 37 | mi_kraskov_NN-4 | Mutual information | Information theory |
| 38 | mi_kraskov_NN-4_DCE | Mutual information | Information theory |
| 39 | si_kernel_W-0.5_k-1 | Stochastic interaction | Information theory |
| 40 | tlmi_gaussian | Time-lagged mutual information | Information theory |

| | | | |
|---|---|---|---|
| 41 | tlmi_kernel_W-0.25 | Time-lagged mutual information | Information theory |
| 42 | tlmi_kraskov_NN-4 | Time-lagged mutual information | Information theory |
| 43 | tlmi_kraskov_NN-4_DCE | Time-lagged mutual information | Information theory |
| 44 | cohmag_multitaper_mean_fs-1_fmin-0_fmax-0-25 | Spectral coherence magnitude | Spectral |
| 45 | plv_multitaper_mean_fs-1_fmin-0_fmax-0-25 | Phase locking value | Spectral |
| 46 | lmfit_BayesianRidge | Linear model fit | Miscellaneous |
| 47 | lmfit_ElasticNet | Linear model fit | Miscellaneous |
| 48 | lmfit_Ridge | Linear model fit | Miscellaneous |
| 49 | lmfit_SGDRegressor | Linear model fit | Miscellaneous |

### 2. Results

#### 2.1 Feature similarity could be used to classify the task labels

Feature similarity showed greater variation across tasks than functional connectivity perhaps because the feature similarity was noisier or more sensitive to the task modulation than functional connectivity. To distinguish these two possibilities, we performed classification analyses to predict task labels for non-semantic tasks (i.e., spatial working memory task versus math task) and semantic tasks (i.e., semantic feature matching versus semantic association task) using feature similarity matrix of each participant, respectively (see Method S.1.2.2 for classification analysis). If the feature similarity matrices could be used to correctly classify the task labels, it suggests that feature similarity carries meaningful task information, and the difference in correlations across tasks between the two measures cannot be attributed solely to noise in feature similarity. We found that the classification accuracies using feature similarity were significantly greater than the chance level for non-semantic tasks (classification accuracy = 0.83, p = 0, chance level = 0.5) and semantic tasks (accuracy = 0.78, p = 0, chance level = 0.5), indicating that feature similarity carries meaningful task information. Functional connectivity could also be used to correctly classify the task labels for non-semantic tasks (classification accuracy = 0.87, p = 0, chance level = 0.5) and semantic tasks (classification accuracy = 0.79, p = 0, chance level = 0.5). These findings indicate that feature similarity captures different aspects of brain function compared to functional connectivity and may provide complementary information in characterizing task-related neural activity.

**2.2 Feature similarity could be used to classify the network labels of parcels**

To reveal network similarity, multi-class classification was used to predict the network labels of parcels using extracted features of the time-series data. These features include temporal autocorrelation, kurtosis, and entropy [44], among others, which may capture meaningful differences between different types of time series and thus represent promising candidates as quantitative phenotypes for distinguishing data of different types. We found that classification accuracy was significantly greater than chance for each participant on each task (rest: mean = 0.37, SD = 0.04; spatial working memory: mean = 0.24, SD = 0.03; math: mean = 0.21, SD = 0.04; semantic feature matching: mean = 0.38, SD = 0.03; semantic association: mean = 0.37, SD = 0.03).

We extracted features of the time-series data for each parcel and conducted a multi-class classification to determine how accurately network labels could be predicted based on parcel features. Linear support vector machine classifiers were trained to predict each parcel's network using its feature vectors. We then analyzed classification outputs for DAN-A and DAN-B. Results showed that DAN-A had a stronger similarity to Visual-A network than DAN-B (Fig. S1). Specifically, DAN-A was most frequently misclassified as Visual-A and FPCN-A, with probabilities of misclassification as Visual-A significantly above chance in all conditions (rest: $t = 25.81$, $p = 5.25e-71$; spatial working memory: $t = 8.76$, $p = 2.25e-9$; math: $t = 8.95$, $p = 1.8e-9$; feature matching: $t = 11.59$, $p = 3.33e-12$; semantic association: $t = 10.34$, $p = 3.46e-11$; FDR-corrected). In contrast, DAN-B's top misclassifications were as Motor-A and DAN-A, with significant Visual-A misclassification probabilities only at rest and in non-semantic tasks (rest: $t = 6.62$, $p = 1.12e-9$; spatial working memory: $t = 4.95$, $p = 8.67e-5$; math: $t = 3.70$, $p = 0.002$; feature matching: $t = 1.55$, $p = 0.165$; semantic association: $t = 1.007$, $p = 0.322$; FDR-corrected). Overall, DAN-A was more likely than DAN-B to be misclassified as Visual-A across rest and all tasks (rest: $t = 14.24$, $p = 4.97e-38$; spatial working memory: $t = 4.19$, $p = 0.0001$; math: $t = 5.59$, $p = 9.61e-7$; feature matching: $t = 8.29$, $p = 3.9e-11$; semantic association: $t = 7.38$, $p = 9.48*e-10$). These findings indicate that DAN-A shares greater similarity with Visual network than DAN-B.

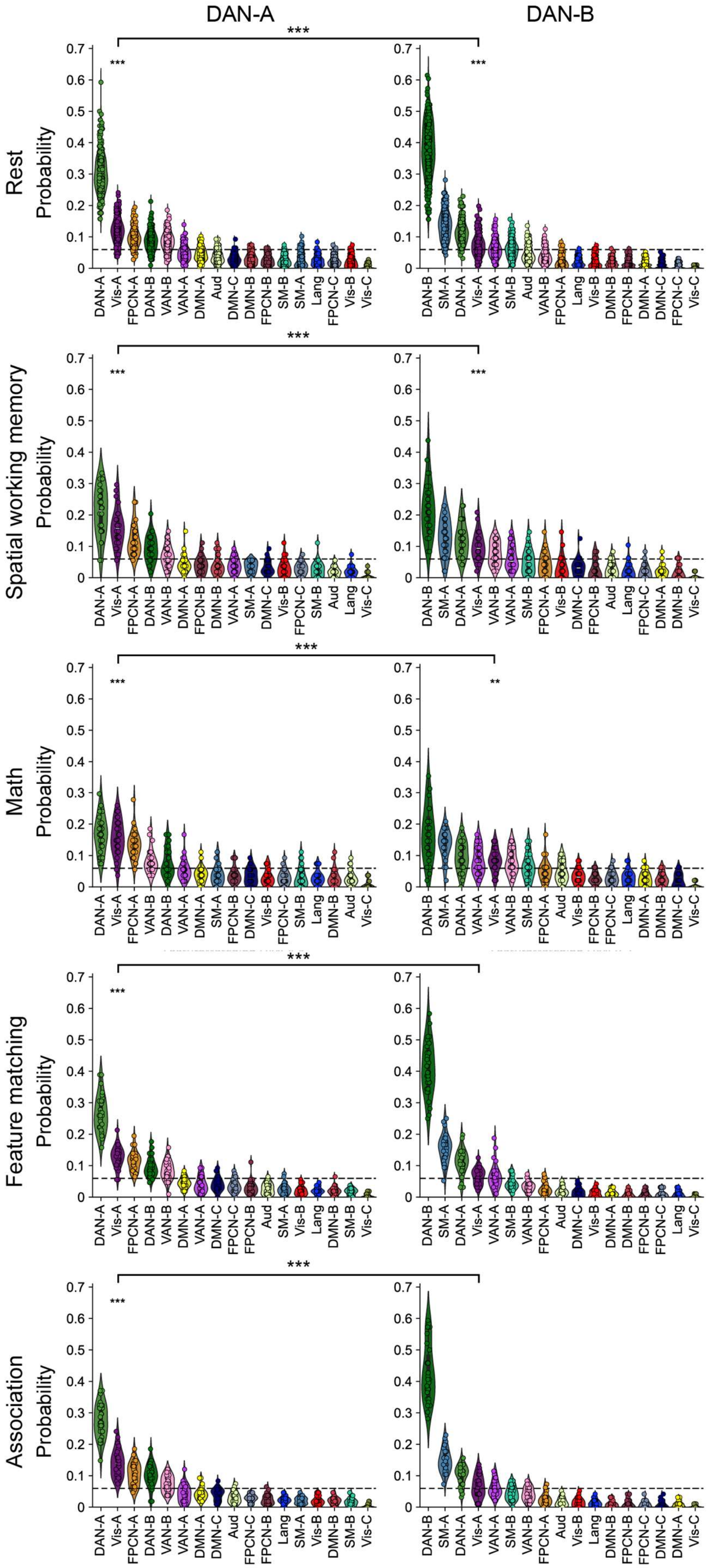
DAN-A
DAN-B
***
**
Rest
Spatial working memory
Math
Feature matching
Association
Probability
0
0.1
0.2
0.3
0.4
0.5
0.6
0.7
DAN-A
DAN-B
Vis-A
Vis-B
Vis-C
FPCN-A
FPCN-B
FPCN-C
VAN-A
VAN-B
DMN-A
DMN-B
DMN-C
SM-A
SM-B
Aud
Lang

Fig. S1. The probabilities of classifying DAN-A and DAN-B as each network. Left panel: The DAN-A was most likely to be misclassified as Visual-A at rest and during tasks. Right panel: The DAN-B was most likely to be misclassified as Motor-A network at rest and during tasks. DAN-A was more likely to be misclassified as Visual-A than DAN-B at rest and during tasks. Dashed lines represent the chance level. **(**Vis = Visual, Aud = Auditory, SM = Sensory-motor, DAN = Dorsal attention network, VAN = Ventral attention network, FPCN = Fronto-parietal control network, Lang = Language, DMN = Default mode network**).**

2.3. FS captured network interaction patterns across tasks not captured by FC

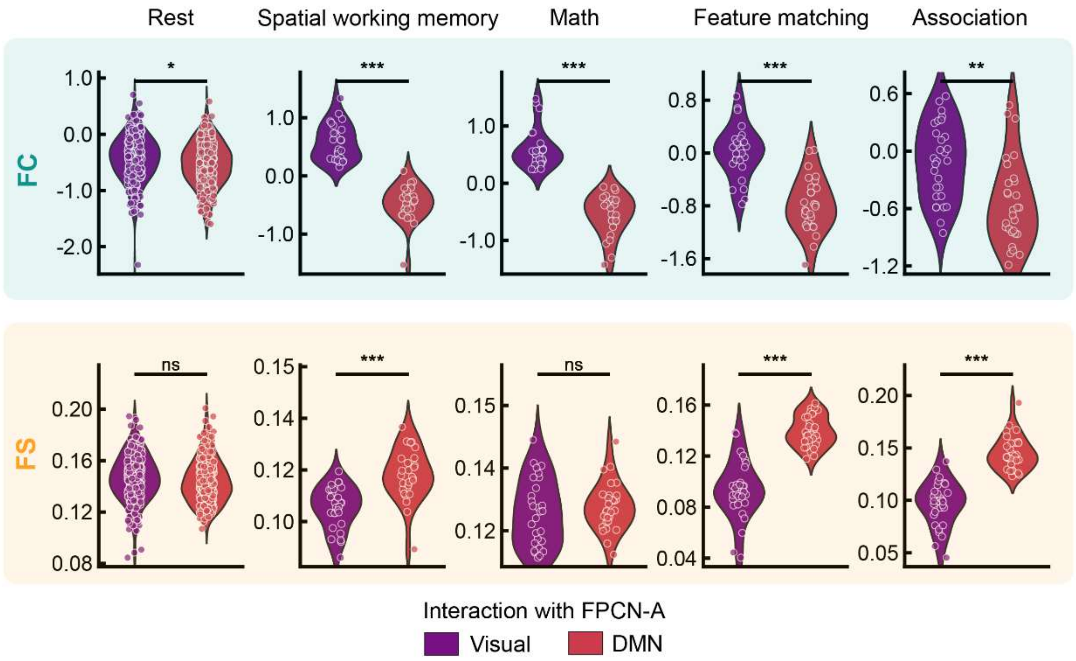


Fig. S2. FS captured the varying interaction patterns across tasks that undetected by FC for FPCN-A. Top panel: FPCN-A always showed greater FC with Visual network than with DMN at rest and for each task and there was no difference across tasks. Bottom panel: FPCN-A showed similar FS with visual and DMN at rest and in the math task but showed greater FS with DMN than with visual network in the spatial working memory task and semantic tasks.

**Table 2. Statistical Results of 49 SPIs for Each Task**

| ID | Statistics | Spatial working memory | | Math | | Feature matching | | Association | |
|---|---|---|---|---|---|---|---|---|---|
| | | t | q | t | q | t | q | t | q |
| 1 | cov_EllipticEnvelope | 11.38 | 0.00 | 8.32 | 0.00 | 9.60 | 0.00 | 7.68 | 0.00 |
| 2 | cov_EmpiricalCovariance | 16.71 | 0.00 | 11.69 | 0.00 | 17.42 | 0.00 | 15.94 | 0.00 |
| 3 | cov_GraphicalLassoCV | 10.27 | 0.00 | 6.24 | 0.00 | 3.82 | 0.00 | 4.26 | 0.00 |
| 4 | cov_LedoitWolf | 15.80 | 0.00 | 11.24 | 0.00 | 17.31 | 0.00 | 15.86 | 0.00 |
| 5 | cov_MinCovDet | 11.38 | 0.00 | 8.32 | 0.00 | 9.76 | 0.00 | 7.69 | 0.00 |
| 6 | cov_OAS | 15.88 | 0.00 | 11.28 | 0.00 | 17.32 | 0.00 | 15.84 | 0.00 |
| 7 | cov_ShrunkCovariance | 16.71 | 0.00 | 11.69 | 0.00 | 17.42 | 0.00 | 15.94 | 0.00 |
| 8 | prec_EllipticEnvelope | -3.70 | 0.00 | -4.44 | 0.00 | -1.68 | 0.14 | -0.78 | 0.44 |
| 9 | prec_EmpiricalCovariance | -2.52 | 0.04 | -3.96 | 0.00 | -0.92 | 0.49 | -0.64 | 0.53 |
| 10 | prec_GraphicalLassoCV | -9.62 | 0.00 | -7.08 | 0.00 | -4.06 | 0.00 | -3.44 | 0.00 |
| 11 | prec_LedoitWolf | -13.78 | 0.00 | -13.08 | 0.00 | -13.24 | 0.00 | -14.50 | 0.00 |
| 12 | prec_MinCovDet | -3.70 | 0.00 | -4.44 | 0.00 | -1.59 | 0.16 | -1.37 | 0.18 |
| 13 | prec_OAS | -13.87 | 0.00 | -13.08 | 0.00 | -13.29 | 0.00 | -14.63 | 0.00 |
| 14 | prec_ShrunkCovariance | -16.42 | 0.00 | -19.16 | 0.00 | -15.51 | 0.00 | -17.71 | 0.00 |
| 15 | spearmanr | 17.13 | 0.00 | 11.49 | 0.00 | 18.07 | 0.00 | 16.45 | 0.00 |
| 16 | kendalltau | 17.02 | 0.00 | 11.43 | 0.00 | 18.10 | 0.00 | 16.37 | 0.00 |
| 17 | reci | -7.32 | 0.00 | -2.58 | 0.03 | 0.50 | 0.83 | 0.06 | 0.96 |
| 18 | bary_euclidean_max | 9.87 | 0.00 | 8.45 | 0.00 | 12.10 | 0.00 | 10.41 | 0.00 |
| 19 | dcorr | 7.87 | 0.00 | 2.76 | 0.02 | -1.08 | 0.29 | -1.36 | 0.24 |
| 20 | dcorr_biased | 7.55 | 0.00 | 2.38 | 0.03 | -1.92 | 0.07 | -2.65 | 0.03 |
| 21 | dtw | -5.12 | 0.00 | -5.74 | 0.00 | -0.82 | 0.56 | -0.28 | 0.78 |
| 22 | dtw_constraint-itakura | -6.24 | 0.00 | -5.55 | 0.00 | -2.59 | 0.02 | -2.94 | 0.01 |
| 23 | dtw_constraint-sakoe-chiba | -16.64 | 0.00 | -11.51 | 0.00 | -17.70 | 0.00 | -15.73 | 0.00 |
| 24 | hsic | 7.75 | 0.00 | 3.07 | 0.01 | -1.09 | 0.29 | -1.24 | 0.29 |
| 25 | hsic_biased | 7.52 | 0.00 | 2.67 | 0.02 | -2.06 | 0.05 | -2.52 | 0.02 |
| 26 | lcss | 13.51 | 0.00 | 8.66 | 0.00 | 9.88 | 0.00 | 9.40 | 0.00 |
| 27 | lcss_constraint-sakoe-chiba | 17.88 | 0.00 | 11.07 | 0.00 | 19.48 | 0.00 | 16.13 | 0.00 |
| 28 | softdtw | -8.84 | 0.00 | -7.31 | 0.00 | -1.31 | 0.26 | -1.16 | 0.26 |
| 29 | softdtw_constraint-itakura | -8.84 | 0.00 | -7.31 | 0.00 | -1.31 | 0.26 | -1.16 | 0.26 |
| 30 | softdtw_constraint-sakoe-chiba | -8.84 | 0.00 | -7.31 | 0.00 | -1.31 | 0.26 | -1.16 | 0.26 |
| 31 | cce_kernel_W-0.5 | -9.85 | 0.00 | -4.11 | 0.00 | -8.73 | 0.00 | -8.62 | 0.00 |

| | | | | | | | | | |
|---|---|---|---|---|---|---|---|---|---|
| 32 | cce_gaussian | 0.39 | 0.70 | 1.70 | 0.13 | -8.34 | 0.00 | -6.20 | 0.00 |
| 33 | ce_gaussian | -8.26 | 0.00 | -3.05 | 0.01 | 0.00 | 1.00 | -0.40 | 0.92 |
| 34 | ce_kernel_W-0.5 | -9.78 | 0.00 | -3.96 | 0.00 | -0.76 | 0.45 | -1.67 | 0.14 |
| 35 | je_gaussian | -8.56 | 0.00 | -3.38 | 0.00 | -0.62 | 0.54 | -2.08 | 0.06 |
| 36 | mi_gaussian | 8.26 | 0.00 | 3.05 | 0.01 | 0.00 | 1.00 | 0.40 | 0.92 |
| 37 | mi_kraskov_NN-4 | 8.14 | 0.00 | 3.14 | 0.01 | -1.12 | 0.34 | -0.97 | 0.34 |
| 38 | mi_kraskov_NN-4_DCE | 7.88 | 0.00 | 3.10 | 0.01 | -0.08 | 0.94 | 0.12 | 0.94 |
| 39 | si_kernel_W-0.5_k-1 | -2.57 | 0.02 | -2.63 | 0.02 | 12.42 | 0.00 | 8.50 | 0.00 |
| 40 | tlmi_gaussian | 6.89 | 0.00 | 2.94 | 0.01 | -0.64 | 0.53 | -1.12 | 0.36 |
| 41 | tlmi_kernel_W-0.25 | 9.15 | 0.00 | 3.15 | 0.01 | 1.28 | 0.21 | 1.49 | 0.20 |
| 42 | tlmi_kraskov_NN-4 | 5.97 | 0.00 | 2.93 | 0.01 | -1.92 | 0.07 | -2.91 | 0.01 |
| 43 | tlmi_kraskov_NN-4_DCE | 6.14 | 0.00 | 2.70 | 0.02 | -0.83 | 0.41 | -1.52 | 0.18 |
| 44 | cohmag_multitaper_mean_fs-1_fmin-0_fmax-0-25 | 8.66 | 0.00 | 2.29 | 0.03 | 11.60 | 0.00 | 11.59 | 0.00 |
| 45 | plv_multitaper_mean_fs-1_fmin-0_fmax-0-25 | 11.49 | 0.00 | 3.34 | 0.00 | 12.96 | 0.00 | 10.72 | 0.00 |
| 46 | lmfit_BayesianRidge | -8.05 | 0.00 | -2.59 | 0.03 | 0.68 | 0.50 | 0.90 | 0.50 |
| 47 | lmfit_ElasticNet | -7.76 | 0.00 | -4.06 | 0.00 | -4.48 | 0.00 | -5.19 | 0.00 |
| 48 | lmfit_Ridge | -8.07 | 0.00 | -2.61 | 0.03 | 0.67 | 0.51 | 0.88 | 0.51 |
| 49 | lmfit_SGDRegressor | -8.07 | 0.00 | -2.61 | 0.03 | 0.67 | 0.51 | 0.88 | 0.51 |

Note. Positive t-values indicate that the DAN-A network exhibits stronger interactions with the Visual network, while negative t-values suggest greater interaction with the DMN. The q-value represents the FDR-corrected p-value.

[15] Lanczos C. Evaluation of Noisy Data. Journal of the Society for Industrial and Applied

Mathematics Series B Numerical Analysis 1964;1:76–85. https://doi.org/10.1137/0701007.

[41] Kelleher JD, Mac Namee B, D'arcy A. Fundamentals of machine learning for predictive data analytics: algorithms, worked examples, and case studies. MIT press; 2015.

[42] Hughes G. On the mean accuracy of statistical pattern recognizers. IEEE Trans Inf Theory 1968;14:55–63. https://doi.org/10.1109/TIT.1968.1054102.

[43] Cliff OM, Bryant AG, Lizier JT, Tsuchiya N, Fulcher BD. Unifying pairwise interactions in complex dynamics. Nat Comput Sci 2023;3:883–93. https://doi.org/10.1038/s43588-023-00519-x.

[44] Shafiei G, Markello RD, De Wael RV, Bernhardt BC, Fulcher BD, Misic B. Topographic gradients of intrinsic dynamics across neocortex. Elife 2020;9:1–24. https://doi.org/10.7554/ELIFE.62116.
